# Active maintenance of accessible chromatin at poised meiotic promoters by the NFYA-NURF axis is required for meiotic entry in mice

**DOI:** 10.64898/2026.10.07.757370

**Authors:** Atiyeh Eghbali, Martin Säflund, Masomeh Askari, Mukhtar Mohamed Abdi, Ann-Kristin Östlund Farrants, Tianxiong Yu, Deniz M. Özata

**Affiliations:** Department of Molecular Biosciences, The Wenner-Gren Institute, Stockholm University, S-106 91 Stockholm, Sweden; Department of Genomics and Computational Biology, University of Massachusetts Medical School, Worcester, MA 01605, USA

**Keywords:** spermatogenesis, NFYA, SMARCA5, BPTF, NURF, chromatin-remodeling factors, chromatin accessibility

## Abstract

Developmental genes often acquire accessible chromatin prior to their activation, yet the mechanisms regulating this permissive state remain poorly understood. During spermatogenesis, many meiotic gene promoters display this poised state in spermatogonia, characterized by accessible promoters and paused RNA Pol II occupancy before their activation during meiosis. Here, we identify the NURF chromatin remodeling complex, comprised of SMARCA5 and its accessory subunit BPTF, as a key regulator of this process. We demonstrate that the transcription factor NFYA interacts with SMARCA5 and is required for its occupancy at poised meiotic gene promoters. Approximately one-third of poised meiotic gene promoters is co-occupied by NFYA, SMARCA5, and BPTF. Germline deletion of *Smarca5* disrupts chromatin accessibility at poised meiotic promoters, impairs transcriptional program required for mitosis-to-meiosis transition, resulting in developmental arrest at meiotic entry. These findings identify NFYA-NURF as a key regulatory axis that regulates accessible chromatin at developmentally poised meiotic genes and primes them for activation later during meiosis.

## Introduction

Spermatogenesis is a stepwise developmental program in which spermatogonia (SpG) undergo mitotic amplification, differentiation, and subsequently enter meiosis^1–3^. Meiosis, a specialized cell division exclusive to germ cells, produces four genetically distinct haploid spermatids that ultimately differentiate into spermatozoa^1–3^. Meiosis represents a highly complex stage of spermatogenesis characterized by programmed DNA double-strand breaks (DSBs), homologous chromosome pairing, and genetic recombination^1–3^. Consequently, meiotic entry and progression are tightly regulated to ensure accurate germ cell development. Disruption of these processes frequently leads to male infertility^1–3^.

The retinoic acid-responsive master transcription factors (TFs) STRA8 and MEIOSIN activate the transcriptional program required for meiotic entry^4–7^, whereas the TFs A-MYB and TCFL5 subsequently promote the expression of genes required for meiotic progression and spermatid differentiation^8–13^. The initiation and progression of meiosis are in fact accompanied by extensive changes in the chromatin landscape^14–20^. Prior to the activity of these master TFs, many meiotic regulatory elements, including promoters and enhancers, already exhibit accessible chromatin and remain poised in pre-meiotic SpG for subsequent activation during meiotic initiation and progression^8,21–23^.

We recently demonstrated that in SpG, the TF NFYA regulates chromatin accessibility at the poised promoters of meiotic genes that become transcriptionally active upon meiotic entry^22^. Previous studies have shown that NFYA can bind to its DNA motif within heterochromatin^24^ and facilitates chromatin accessibility at regulatory elements that are subsequently bound by master TFs in mouse embryonic stem cells^25^. Collectively, these findings, together with our recent work, supports a model in which NFYA functions as a pioneer factor to establish and/or maintain permissive chromatin at poised meiotic promoters in SpG. One of the salient properties of pioneer factors is to recruit nucleosome-remodelling complexes to promote accessible chromatin^26–29^.

Here, we demonstrate that SMARCA5 (SNF2H), an ATPase subunit of the ISWI chromatin-remodeling complex, and its accessory subunit BPTF – together, they form the NURF complex – interact with NFYA. Notably, NFYA and the NURF complex co-occupy approximately one-third of the poised meiotic promoters in SpG, including those activated by STRA8 and MEIOSIN. SMARCA5 occupancy at these poised promoters is markedly reduced in NFYA-deficient SpG, suggesting that NFYA recruits the NURF complex to such promoters prior to meiotic entry. Spermatogenesis arrests at meiotic entry in mice with a germline-specific deletion of *Smarca5*. Finally, poised promoters occupied by the NURF complex in the absence of NFYA binding are instead bound by the pioneer factors ZBTB16 and SALL4. Together, our work uncovers a key regulatory role for the NFYA-SMARCA5-BPTF axis in facilitating chromatin accessibility at poised meiotic promoters and its intricate interplay with other pioneer factors.

## Results

### NFYA recruits the ISWI ortholog SMARCA5 to poised meiotic gene promoters in spermatogonia

We recently identified NFYA as a regulator of accessible chromatin at poised meiotic gene promoters prior to meoisis^22^ attributing to its potential pioneer activity. One of the hallmarks of pioneer factors is to recruit chromatin-remodeling factors^26–29^. To characterize NFYA-interacting proteins, we immunoprecipitated endogenous NFYA from the testis and liver tissues of adult mice using antibodies against NFYA and IgG and performed mass spectrometry (MS) from the immunoprecipitates (Figures 1A, 1B, S1A and Table S1). To catalogue testis-specific interacting partners, liver served as a control tissue. MS analysis revealed that SMARCA5 and BPTF enriched over IgG control specifically in the testes of adult mice (Figures 1A, 1B, S1A). SMARCA5 is an ATPase of imitation switch (ISWI) remodeler; association of SMARCA5 with BPTF – an ISWI accessory subunit – forms the NURF complex^30–32^.

**Figure 1.**
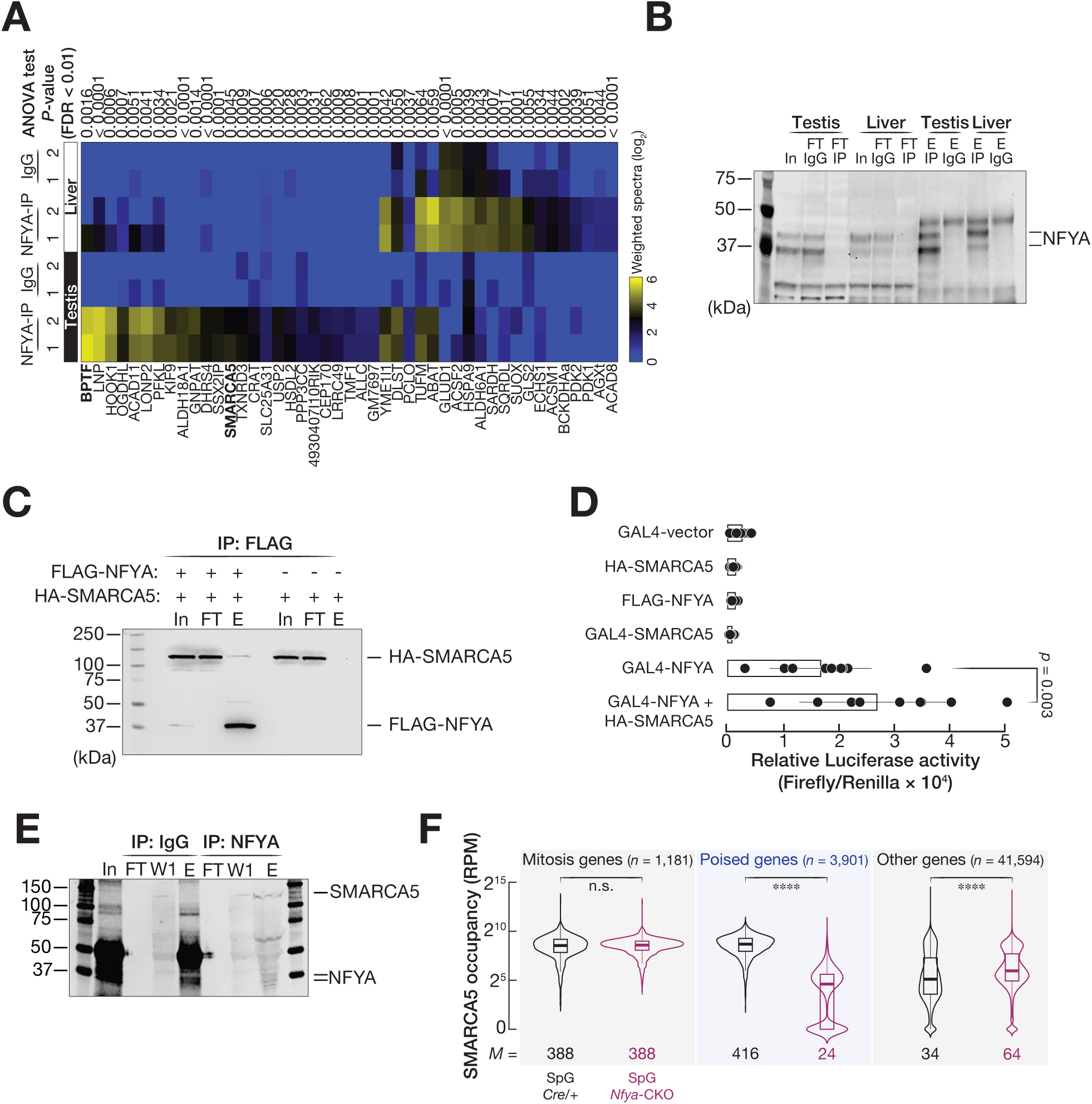
NFYA interacts with SMARCA5 and is required for SMARCA5 occupancy at poised meiotic gene promoters in spermatogonia. See also Figure S1. (A) Heatmap shows log_₂_-transformed weighted spectral values of proteins identified by mass spectrometry following immunoprecipitation of endogenous NFYA from adult mouse testis and liver extracts. IgG immunoprecipitations served as negative controls, and liver served as a control tissue to identify testis-enriched NFYA-interacting proteins. Two independent replicates are shown. ANOVA test is applied to measure the significant differences between conditions (false discovery rate [FDR] < 0.01). (B) Western blotting shows the abundance of NFYA protein in the eluate fractions from NFYA and IgG immunoprecipitations using adult mouse testis and liver protein extracts. In, input; FT, flow-through; E, eluate. (C) Western blotting upon co-immunoprecipitation of FLAG-tagged NFYA and HA-tagged SMARCA5 in HEK293T cells shows the abundances of FLAG-tagged NFYA and HA-tagged SMARCA5 proteins. Co-immunoprecipitation is performed using anti-FLAG M2 paramagnetic beads. The membrane is blotted with both anti-FLAG and anti-HA antibodies. Cells expressing HA-tagged SMARCA5 alone serves as a negative control. In, input; FT, flow-through; E, eluate. (D) Relative luciferase activity in HEK293T cells stably carrying a UAS-driven firefly luciferase reporter and expressing the indicated plasmid DNA. Firefly luciferase activity was normalized to Renilla luciferase activity. GAL4-vector, HA-SMARCA5, FLAG-NFYA, and GAL4-SMARCA5 plasmids serve as controls. The whiskers show the standard deviation. Significance is measure using two-sided paired Student *t* test. (E) Western blotting shows the abundances of NFYA and SMARCA5 proteins after immunoprecipitation of endogenous NFYA from adult mouse testis extracts. IgG immunoprecipitation served as a negative control. In, input; FT, flow-through; E, eluate. (F) Violin plots show reads per million (RPM)-normalized SMARCA5 CUT&Tag signals at the promoters of mitosis genes (*n* = 1,181), poised genes (*n* = 3,901), and other genes excluding mitotic and poised genes (*n* = 41,594) in FACS-purified spermatogonia (SpG) from male mice with germ cell-specific deletion of endogenous *Nfya* (*Nfya*-*CKO*; magenta) and control *Stra8-Cre^Cre/+^* control males (*Cre/+*; black). \*\*\*\**p* < 0.0001; Mitotic genes in *Cre/+* SpG vs *Nfya*-*CKO p* = n.s., Poised genes in *Cre/+* SpG vs *Nfya*-*CKO p* < 2.2 × 10^-16^, Other genes *Cre/+* SpG vs *Nfya*-*C*KO *p* < 2.2 × 10^-16^; two-sided wilcoxon matched-pairs signed-ranked sum test. Whiskers show maximum and minimum values. Interquartile range (IQR) is represented by boxplots.

To validate the physical interaction between SMARCA5 and NFYA, we ectopically co-expressed FLAG-tagged NFYA (FLAG-NFYA) and HA-tagged SMARCA5 (HA-SMARCA5) in heterologous HEK293T cells and performed immunoprecipitation (IP) using anti-FLAG antibody. IP followed by western blotting revealed that HA-SMARCA5 co-precipitated with FLAG-NFYA (Figures 1C, S1B). To assess whether the physical interaction between NFYA and SMARCA5 has functional consequence on transcriptional activation, we co-expressed GAL4-tagged NFYA (GAL4-NFYA) along with HA-SMARCA5 in HEK293T cells that stably express firefly luciferase gene under UAS promoter. Ectopic expression of GAL4-NFYA-alone resulted in the activation of firefly luciferase gene, whereas luciferase activity was not detected in cells that either express GAL4-SMARCA5, or FLAG-NFYA, or HA-SMARCA5 (Figure 1D). However, co-expression of GAL4-NFYA and HA-SMARCA5 markedly increased the transcription of firefly luciferase gene when compared to cells expressing only GAL4-NFYA, substantiating the functional relevance of physical interaction between NFYA and SMARCA5 (Figure 1D; two-sided paired Student *t* test; *p* = 0.003).

Concordantly, immunoprecipitation of endogenous NFYA from adult mouse testes followed by western blotting further validated the physical interaction between NFYA and SMARCA5 (Figures 1E, S1C). In our previous work, we demonstrated that NFYA binds to the promoters of both mitotic and poised meiotic genes in SpG^22^. To directly test whether SMARCA5 occupancy in SpG is dependent on NFYA, we performed CUT&Tag for SMARCA5 in FACS-purified SpG from males in which endogenous *Nfya* was specifically deleted in germ cells using STRA8 promoter-driven Cre recombinase (Henceforth, *Nfya*-CKO)^22^. SpG from *Stra8-Cre^Cre/+^* males (Henceforth, *Cre/+*) served as controls. Intriguingly, SMARCA5 occupancy at poised meiotic gene promoters was reduced ∼20-fold in SpG from *Nfya*-CKO compared with that from *Cre/+*, whereas SMARCA5 occupancy at promoters of mitotic genes remained unchanged (Figure 1F; two-sided wilcoxon matched-pairs signed-ranked sum test; *p* < 0.001). Together, these findings demonstrate that NFYA recruits SMARCA5 to the promoters of poised meiotic genes in SpG.

### The expression of SMARCA5 is enriched in pre-meiotic germ cells

NFYA is abundantly expressed in SpG and pre-leptotene (pL)^22^. We sought to determine whether the spatiotemporal expressions of SMARCA5 and BPTF align with that of NFYA during spermatogenesis. Analysis of our publicly available single-cell RNA sequencing (scRNA-seq) data^22^ revealed that, similar to *Nfya*, the mRNA abundances of *Smarca5* and *Bptf* were higher in pre-meiotic germ cells, SpG and pL, than in other developing meiocytes and spermatids (Figure S2A). We next performed SMARCA5 immunofluorescence staining on seminiferous tubules from the testes of staged mice. At 9 days post-partum (dpp), when developing SpG constitute the predominant germ cell population and meiosis I has only recently sterted, progressing no further than leptotene (L)^33^, SpG exhibited strong nuclear SMARCA5 staining (Figure S2B). At 14 dpp, when meiosis I reaches early pachytene (eP) stage^33^, SpG located along the basal membrane showed strong nuclear SMARCA5 signal, whereas eP lacked detectable nuclear SMARCA5 signal (Figure S2B). By 17 to 21 dpp, when meiosis is complete and spermatids are readily apparent^33^, developing SpG displayed stronger nuclear SMARCA5 signal than meiocytes and spermatids (Figure S2B). Note that the specificity of the SMARCA5 antibody was tested using testes from *Smarca5* mutant mice (Figure S3). Taken together, we conclude that SMARCA5 is predominantly expressed in pre-meiotic germ cells.

### SMARCA5-deficient male mice are sterile

Given that both NFYA and the NURF complex are highly expressed in pre-meiotic cells^22^ (Figures 1, S1, S2) and that NFYA is essential for meiotic initiation^22^, we sought to address the biological role of SMARCA5 in meiotic entry and progression. To this end, we generated *Smarca5^fl/fl^*; *Stra8-Cre* mice in C57BL/6J background, in which exon 5 of *Smarca5* is specifically deleted in germ cells (Figures S3A-C; henceforth, *Smarca5-CKO*). The *Stra8-Cre* mice, generated in C57BL/6J background by Cyagen, harbor a *P2A-ZsGreen1-T2A-Cre* cassette inserted immediately upstream of *Stra8* stop codon, resulting in Cre recombinase expression in pre-meiotic cells (Figure S3A; “STAR Methods”). Western blotting confirmed depletion of SMARCA5 in the testes of 14 and 30 dpp mice, while immunofluorescence staining validated the loss of SMARCA5 specifically in germ cells (Figures S3C, D). Consistently, immunostaining for the Sertoli cell marker SOX9^34^ revealed comparable number of SOX9-positive cells in *Smarca5-CKO* and *Cre/+* mice at both 14 and 30 dpp (Figure S3E). *Smarca5-CKO* males had significantly smaller testes than *Cre/+* (Figures S4A, B). Furthermore, histological analysis of caudal epididymides from adult *Smarca5-CKO* males revealed complete absence of mature spermatozoa (Figure S4C). We thus conclude that *Smarca5-CKO* mice are sterile.

### SMARCA5 is required for meiotic initiation

We performed hematoxylin and eosin (H&E) staining on the testis sections collected at 14 and 30 dpp. Histological examination revealed severe impairment of spermatogenesis in *Smarca5-CKO* mice (Figure 2A). In *Cre/+* mice, spermatogenesis had progressed to eP stage of meiosis I by 14 dpp, whereas spermatids became readily apparent at 30 dpp (Figure 2A). In contrast, in the tubules of *Smarca5-CKO* mice, the most advanced germ cells morphologically resembled pL spermatocytes (Figure 2A; pL-like spermatocytes). Consistent with the impaired spermatogenesis in *Smarca5-CKO* mice, immunostaining for DDX4, a germ cell marker^35^, revealed a marked reduction in the number of DDX4-positive germ cells at both 14 and 30 dpp (Figure S5A). Furthermore, terminal dUTP nick-end labeling (TUNEL) staining revealed significantly higher number of TUNEL-positive germ cells in the testes of *Smarca5-CKO* mice at 14 and 30 dpp compared with *Cre/+* (Figure S5B), suggesting that defective germ cells are eliminated by apoptosis. We subsequently performed immunostaining for HSPA2, a marker for meiocytes and spermatids^36^, and peanut agglutinin (PNA), a marker for spermatids^37^. We found that HSPA2- and PNA-positive germ cells were absent in *Smarca5-CKO* mice (Figures S5C, D), further substantiating that the most advanced germ cells in *Smarca5-CKO* mice are pL-like spermatocytes that fail to progress through meiotic prophase I.

**Figure 2.**
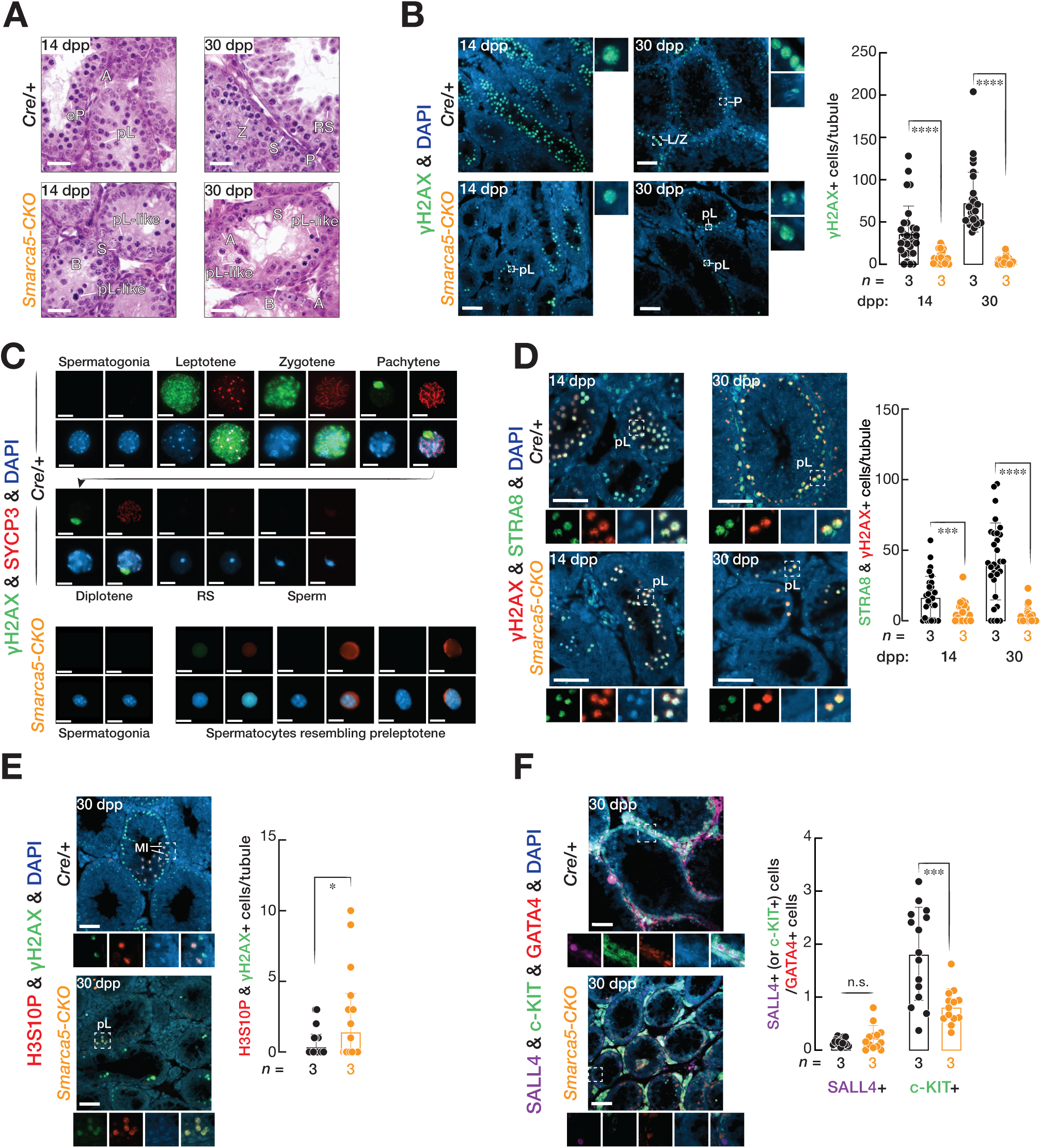
SMARCA5 is essential for meiotic entry and progression. See also Figures S2, S3, S4, S5. **(A)** Hematoxylin and eosin staining of testis sections from *Cre/+* and *Smarca5-CKO* mice at 14 and 30 days post-partum (dpp). A, type A spermatogonia (SpG); B, type B SpG; pL-like, preleptotene-like spermatocytes; Z, zygotene; eP, early pachytene; P, pachytene; RS, round spermatid; S, Sertoli cell. Scale bars: 50 μm. **(B)** Immunofluorescence staining for γH2AX in testis sections from *Cre/+* and *Smarca5-CKO* mice (*n* = 3) at 14 and 30 dpp. Insets show enlarged views of the boxed regions. Images are representative of three biological replicates. Bars represent quantification of γH2AX-positive cells per seminiferous tubule from three biological replicates. Whiskers show standard deviation and the center of the whiskers represents the mean. \*\*\*\**p* < 0.0001; 14 dpp, *Cre/+* vs *Smarca5-CKO p* = 9.4 × 10^-6^; 30 dpp, *Cre/+* vs *Smarca5-CKO p* = 9.2 × 10^-16^; two-sided unpaired Student *t* test. Scale bars: 50 μm. **(C)** Immunofluorescence staining for γH2AX and SYCP3 on chromosome spreads of 8-week-old *Cre/+* and *Smarca5-CKO* testes. Scale bars: 10 μm. **(D–F)** Immunofluorescence staining of the testis sections from *Cre/+* and *Smarca5-CKO* mice at 14 dpp and 30 dpp for **(d)** γH2AX and STRA8 (14 dpp, *n* = 3, *p* = 0.0003; 30 dpp; *n* = 3, *p* = 1.5 × 10^-10^), **(e)** H3S10P and γH2AX (30 dpp; *n* = 3, *p* = 0.04; MI, first meiotic division; pL, preleptotene), **(f)** SALL4, c-KIT, and GATA4 (30 dpp; *n* = 3; SALL4-positive cells, *p* = n.s.; c-KIT-positive cells, *p* = 0.0007). Insets show enlarged views of the boxed regions. Images are representative of three biological replicates. Bars represent quantification for positive cells per tubule from three biological replicates. Whiskers show standard deviation, and the center of the whiskers represents the mean. Significance is measured by two-sided unpaired Student *t* test. Scale bars: 50 μm.

The spermatogenic defect in *Smarca5-CKO* mice appeared to occur at meiotic entry judged by the morphology of pL-like spermatocytes (Figure 2A). Indeed, the observed phenotype in *Smarca5-CKO* mice closely resembled that in *Nfya-CKO* mice, in which NFYA-deficient pL spermatocytes fail to progress through meiosis I^22^. The meiotic entry and progression are characterized by the formation of programmed DNA double-strand breaks (DSB) and the timely expression of proteins required for homologous recombination^38,39^. Immunostaining for γH2AX, a marker for DBS^2^, demonstrated a significantly higher number of spermatocytes with DSBs in *Cre/+* testis at both 14 and 30 dpp (Figure 2B). Notably, however, prominent γH2AX signals were also detected in the nuclei of pL-like spermatocytes of *Smarca5-CKO* mice (Figure 2B). To further characterize these cells, we performed double staining for γH2AX and SYCP3 on the chromosome spreads prepared from *Cre/+* and *Smarca5-CKO* mice. While the full trajectory of germ cells was observed in *Cre/+* mice, *Smarca5-CKO* testes contained only SpG and pL-like spermatocytes displaying diffused SYCP3 signal (Figure 2C). The presence of diffusive SYCP3-positive pL-like spermatocytes indicates defective synaptonemal complex formation and meiotic progression.

STRA8 is essential for meiotic entry and is highly expressed in pL^4,7,40–43^. To determine whether SMARCA5 is required for meiotic entry, we performed double staining for STRA8 and γH2AX on the testis sections from *Cre/+* and *Smarca5-CKO* mice at 14 and 30 dpp. Notably, many γH2AX-positive pL-like spermatocytes exhibited strong nuclear STRA8 staining (Figure 2D). In *Nfya-CKO* males, pL spermatocytes failed to transition from mitosis to meiosis and instead retain a precocious mitotic status^22^. We therefore examined whether SMARCA5-deficient pL-like spermatocytes display a similar phenotype. Double immunostaining for H3S10P (histone H3 Ser10 phosphorylation), a marker for mitotic chromosome condensation^44^, and γH2AX revealed that many γH2AX-positive pL-like spermatocytes in *Smarca5-CKO* mice exhibited nuclear H3S10P staining. In contrast, in *Cre/+* mice, H3S10P staining was largely confined to spermatocytes undergoing metaphase I (Figure 2E). Taken together, these findings demonstrate that, similar to *Nfya-CKO* mice, SMARCA5-deficient germ cells fail to properly transition from mitosis to meiosis and consequently arrest at meiotic entry.

### SMARCA5-deficient male mice exhibit aberrant spermatogonial differentiation

SMARCA5 is expressed abundantly prior to meiotic entry (Figure S2). However, in *Smarca5-CKO* mice, spermatogenesis progress no further than pL, which is the immediate descendants of differentiating SpG. To determine whether SMARCA5 is required for the maintenance and differentiation of SpG, we performed triple immunostaining for SALL4, a marker of undifferentiated SpG^45^; c-KIT, a marker of differentiating SpG^46^; and GATA4, a Sertoli cell marker^47,48^. Because Sertoli cells were unaffected in *Smarca5-CKO* males (Figure S3E), the numbers of SALL4-positive or c-KIT-positive SpG were normalized to the number of GATA4-positive Sertoli cells. The number of SALL4-positive undifferentiated SpG was comparable between *Cre/+* and *Smarca5-CKO* mice (Figure 2F). In contrast, the number of c-KIT-positive differentiating SpG was reduced by ∼2-fold in *Smarca5-CKO* compared to *Cre/+* (Figure 2F). These results indicate that although the maintenance of undifferentiated SpG is unaffected in *Smarca5-CKO* testes, deletion of *Smarca5* impairs spermatogonial differentiation.

### The NURF complex is involved in regulating the mitotic transcriptional program

To investigate the molecular basis of impaired meiotic initiation in *Smarca5-CKO* males, we performed simultaneous scRNA-seq and scATAC-seq on testicular nuclei from 8-week-old *Smarca5-CKO* mouse and integrated these data with our previously published datasets from age-matched *Cre/+* and *Nfya-CKO* mice. The combined scRNA-seq analysis of the three datasets captured 78,860 individual cells, with averages of 2,281 and 5,537 detected genes and transcript counts (unique molecular indices, UMIs) per cell, respectively (Table S2A). UMAP analysis identified 33 transcriptionally distinct cell clusters that were resolved into 15 distinct cell populations using 36 cell-type specific markers, encompassing the full germ cell developmental trajectory (Figures 3A–D and Table S2B). For example, clusters 18 and 30 were identified as undifferentiated spermatogonia (undiff. SpG), based on the enriched expression of *Sall4* and *Zbtb16*, whereas cluster 13 was classified as differentiating spermatogonia (diff. SpG) due to its increased expression of *Stra8* and *Prdm9* (Figure 3E). However, cells from clusters 16 and 24 – originating from *Nfya-CKO* and *Smarca5-CKO*, respectively – exhibited both undiff. and diff. SpG markers and were therefore collectively classified as SpG (Figures 3B–E). Similarly, meiocyte clusters were marked by enriched *Mybl1* and *Piwil1* expression, while spermatid clusters revealed increased *Prm1* and *Sun5* expression (Figure 3E). Genotype-based UMAP visualization revealed the presence of full germ cell population in *Cre/+*, whereas meiocyte populations were absent in both *Nfya-CKO* and *Smarca5-CKO* (Figure 3F). This finding is consistent with the spermatogenic arrest at meiotic entry in *Nfya-CKO*^22^ and *Smarca5-CKO* males (Figure 2).

**Figure 3.**
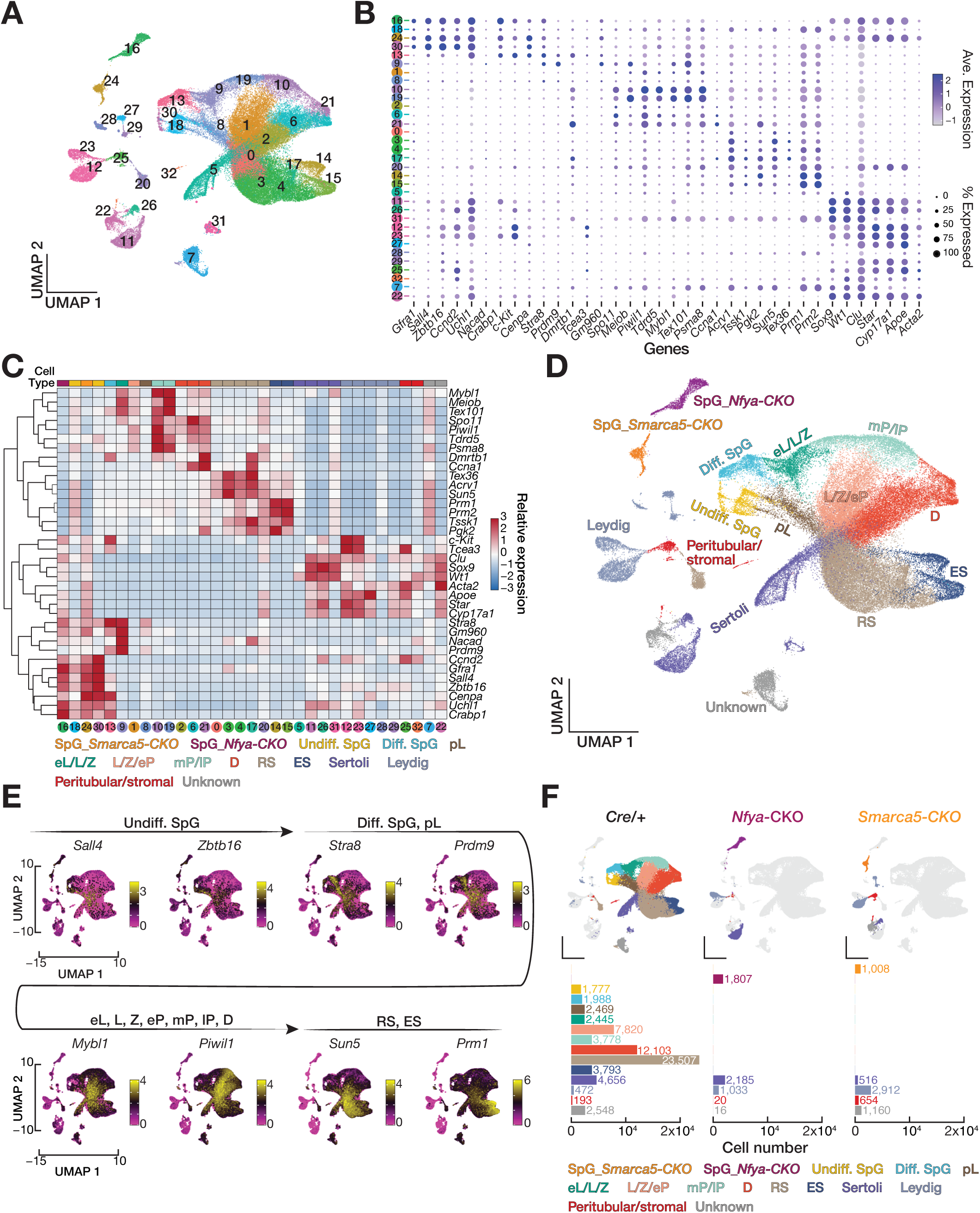
scRNA-seq from reveals lack of meiocytes in testis of *Smarca5-CKO* mouse. **(A)** Dimension reduction analysis by uniform manifold approximation and projection (UMAP) on 78,860 individual cells captured by combined single-cell RNA sequencing (scRNA-seq) datasets from 8-weeks old *Cre/+*, *Smarca5-CKO* and *Nfya-CKO* testes illustrates 32 cell clusters. **(B)** Dot plot shows the average expression (color gradient) and the percentages of cells (dot size) of each cluster for 36 cell-type specific markers. **(C)** Unsupervised hierarchical clustering of 32 cell clusters based on the relative expression levels of 36 marker genes identified 15 cell populations. SpG_*Smarca5-CKO* spermatogonia from *Smarca5-CKO*, SpG_*Nfya-CKO* SpG from Nfya-CKO, Undiff. SpG, undifferentiated SpG; Diff. SpG, differentiating SpG; pL, preleptotene; eL/L/Z, early leptotene, leptotene, and zygotene; L/Z/eP, leptotene, zygotene, and early pachytene; mP/lP, middle and late pachytene; D, diplotene; RS, round spermatid; ES, elongating spermatid; Sertoli cells; Leydig cells; Peritubular/stromal cells; Unknown. **(D)** 15 major cell populations resolved from 32 cell clusters are marked on UMAP space. **(E)** Per-cell gene expression patterns of several exemplified marker genes are visualized on UMAP space. Direction of germ cell developmental trajectory is indicated by black arrows. **(F)** UMAP spaces (top) illustrate the origin of cell populations identified in *Cre/+*, *Smarca5-CKO* and *Nfya-CKO* testes. Bars (bottom) represent the total number of individual cells from each major cell population in each mouse.

*Smarca5* and *Bptf* are abundantly expressed in pre-meiotic germ cells (Figure S2A). Consistently, we found that *Smarca5-CKO* mice exhibit impaired spermatogonial differentiation (Figure 2F) and that SMARCA5-deficient SpG form a distinct cluster from *Cre/+* SpG in UMAP space (Figure 3D), suggesting that the NURF complex regulates mitotic transcriptional program. Using scRNA-seq, we measured changes in transcript abundance in SMARCA5-deficient SpG compared with that in *Cre/+*. In SMARCA5-deficient SpG, 5,607 genes exhibited significantly altered RNA abundance: the abundance of 1,614 genes was decreased, whereas that of 3,993 genes was increased (Figure 4A; *Smarca5-CKO* ÷ *Cre/+* ≤ 0.5 or ≥2; FDR < 0.01; and Table S3). Notably, gene ontology (GO) analysis showed that reduced genes were enriched for male reproductive processes, including meiotic nuclear division, whereas increased genes were associated with gene categories including non-germ cell differentiation and macromolecule metabolism (Figure 4B and Table S4), suggesting that SMARCA5 both positively and negatively regulates mitotic transcriptional program. CUT&Tag for SMARCA5 and BPTF from FACS-purified SpG (Figure S6A and B; agreement between three biological replicates; Pearson correlation coefficient (*r*); SMARCA5 *r* > 0.98; BPTF *r* > 0.85), together with our previously published NFYA CUT&RUN data, revealed that the promoters of approximately 33% of increased genes (1,289 of 3,993) were occupied by SMARCA5 alone, without NFYA and BPTF binding (Figure 4C and Table S3). In addition to BPTF, SMARCA5 can associate with alternative accessory proteins to form distinct remodeling complexes that may mediate gene repression^31,49^. We thus examined the histone modifications associated with gene activation and repression at promoters of genes increased in SMARCA5-deficient SpG, using wild-type diff. SpG as a reference. Intriguingly, our re-analysis of published datasets ^50,51^ for the occupancies of H3K27ac (histone H3 Lys27 acetylation), a modification of active transcription^52^, and H2AK119ub (histone H2 Lys119 ubiquitination), a Polycomb-associated repressive mark^53^, revealed that increased genes with promoters bound by SMARCA5 alone exhibited lower H3K27Ac but higher H2AK119ub signals than increased genes co-occupied by NFYA and the NURF complex (Figure 4D; Mann–Whitney–Wilcoxon *U* test). As SMARCA5 may also indirectly repress genes by suppressing transcriptional activators, we performed HOMER motif enrichment analysis on the promoters of increased genes (Benjamini–Hochberg multiple correction test, FDR < 0.05). The Sp/Klf family binding motif (5’-RGGGYGY-3’) was significantly enriched among these promoters (Figure S7). Notably, the RNA abundances of *Sp5* and *Klf15*, regulators of developmental and metabolic pathways, respectively^54–56^, were increased in SMARCA5-deficient SpG (Figure 4E). In wild-type SpG, their promoters were bound by SMARCA5 alone and displayed low H3K27Ac but high H2AK199ub signal (Figure 4E, two-sided Wilcoxon matched-pairs signed-ranked sum test).

**Figure 4.**
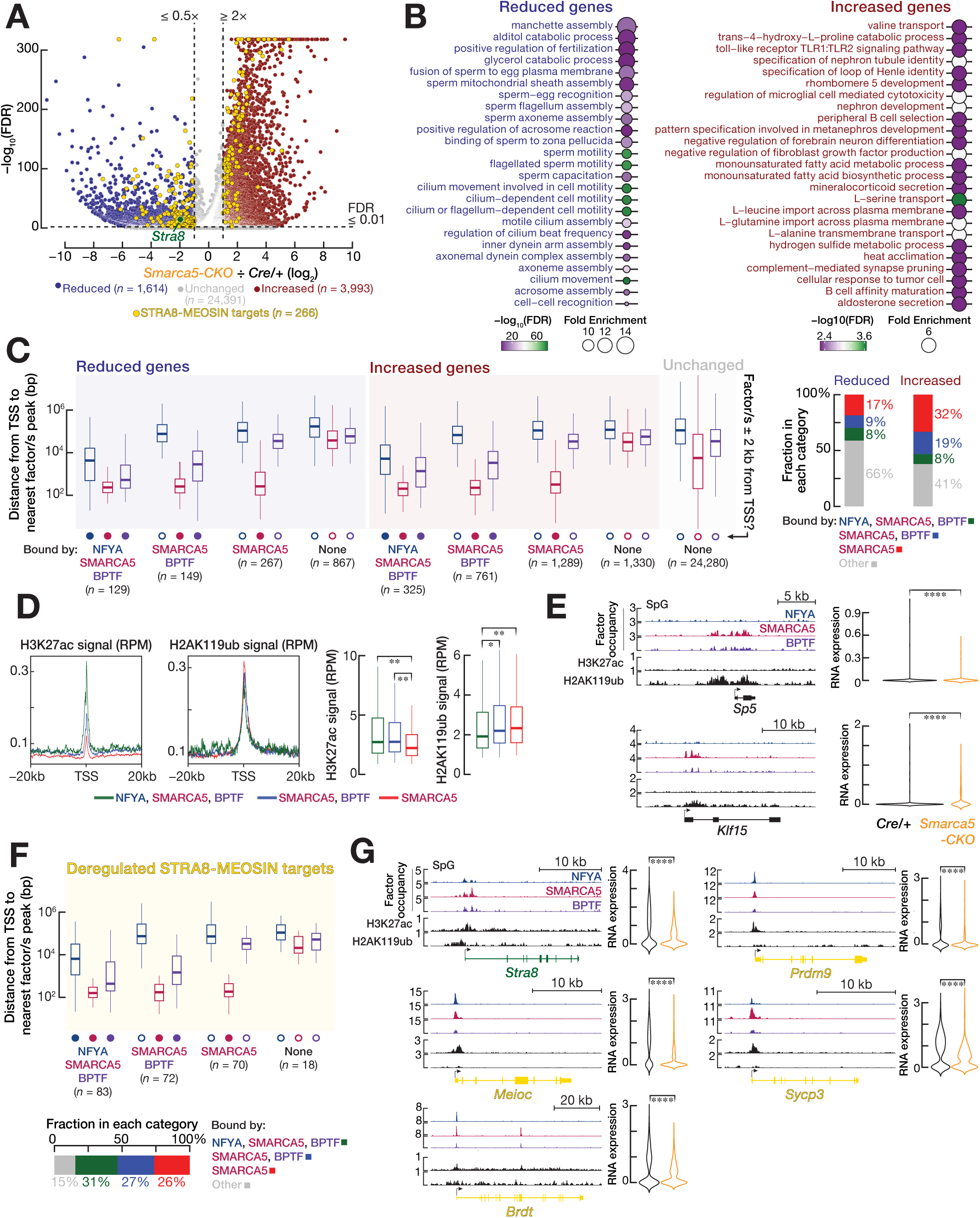
SMARCA5 is required for the transition from mitosis to meiosis. See also Figures S6 and S7. **(A)** Volcano plot shows genes with significantly altered RNA abundance in spermatogonia (SpG) from *Smarca5-CKO* compared to *Cre/+*, determined by scRNA-seq. Genes with significantly reduced RNA abundance in *Smarca5-CKO* SpG (FC ≤ 0.5, FDR ≤ 0.01) are shown in blue, whereas genes with significantly increased RNA abundance in *Smarca5-CKO* SpG (FC ≥ 2, FDR ≤ 0.01) are shown in red. The *Stra8* gene is marked in green. Those deregulated genes, which are controlled by STRA8-MEIOSIN regulatory axis, are highlighted in yellow. Benjamini–Hochberg correction was used to calculate false discovery rate. **(B)** Geno Ontology (GO) analysis for reduced (left) and increased (right) genes. Top 25 gene categories, ranked by fold enrichment, are shown (FDR < 0.01). **(C)** Box plots (left) show the distance (average of three biological replicates) from the transcription start site (TSS) to the nearest NFYA, SMARCA5, or BPTF peak for reduced (*n* = 1,614), increased (*n* = 3,993), or unchanged (*n* = 24,280) genes in SpG. Genes are classified into four categories based on NFYA, SMARCA5, and BPTF occupancy: genes co-bound by all three factors, genes co-bound by SMARCA5 and BPTF, genes bound by SMARCA5 alone, and genes not bound by any of these factors. A Gene was considered as factor-bound if a significant peak was detected within ±2 kb of its TSS in at least two biological replicates. Horizontal lines indicate the median. Whiskers show the minimum and maximum values. Boxes represent the interquartile range (IQR). Bars (right) represent the fraction of each gene category over among all reduced and increased genes. **(D)** Metagene plots (left) show the reads per million (RPM)-normalized H3K27ac and H2AK119ub signals in the –20 kb to +20 kb window flanking TSSs of genes co-bound by all three factors (green), co-bound by SMARCA5 and BPTF (blue), and bound by SMARCA5 alone (red) in wild-type SpG. Box plots (right) show RPM-normalized H3K27ac and H2AK119ub occupancy around the TSSs of each gene category in wild-type SpG. Horizontal lines indicate the median. Boxes represent the IQR. Whiskers show the minimum and maximum values. Statistical significance is measured by Mann–Whitney–Wilcoxon *U* test. **(E)** Integrative Genomics Viewer (IGV) tracks (left) showing NFYA, SMARCA5, BPTF occupancy (RPM), together with H3K27ac and H2AK119ub signals (RPM), at gene boundaries of *Sp5* and *Klf15* in wild-type SpG. Violin plots (right) show RNA abundances of *Sp5* and *Klf15* in SMARCA5-deficient SpG compared with *Cre/+* SpG. Statistical significance is measured by two-sided Wilcoxon matched-pairs signed-ranked sum test. **(F)** Box plots (top) show the distance (average of three biological replicates) from the TSS to the nearest NFYA, SMARCA5, or BPTF peak for deregulated genes (*n* = 266), which are controlled by STRA8-MEIOSIN regulatory axis, in wild-type SpG. Numbers of genes in each category are indicated below the plots. Horizontal lines represent the median, boxes represent the IQR, and whiskers show the minimum and maximum values. Bars (bottom) represent the fraction of each gene category over deregulated STRA8-MEIOSIN target genes. **(G)** IGV tracks (left) showing NFYA, SMARCA5, BPTF occupancy (RPM), together with H3K27ac and H2AK119ub signals (RPM), at the exemplified genes that are controlled by STRA8-MEIOSIN regulatory axis in wild-type SpG: *Stra8*, *Prdm9*, *Meioc*, *Sycp3* and *Brdt*. Violin plots (right) show RNA abundance of each exemplified gene in SMARCA5-deficient SpG compared with *Cre/+* SpG. Statistical significance is measured by two-sided Wilcoxon matched-pairs signed-ranked sum test.

The STRA8–MEIOSIN regulatory axis is essential for meiotic initiation in male germ cells^2,5,6^. Consistent with the spermatogenic defect in meiotic entry observed in *Smarca5-CKO*, SMARCA5-deficient SpG exhibited reduced *Stra8* expression together with altered expression of 265 STRA8–MEIOSIN target genes (Figure 4A). Promoters of 31% of these genes were co-occupied by NFYA, SMARCA5, and BPTF (Figure 4F), indicating that impaired meiotic initiation is at least partly attributable to reduced *Stra8* expression and deregulation of STRA8–MEIOSIN targets. Notably, the subset of STRA8–MEIOSIN target genes that were reduced in *Smarca5-CKO* SpG and whose promoters were co-occupied by NFYA, SMARCA5 and BPTF included key regulators of meiotic recombination, such as *Prdm9* ^39,57–59^ and *Sycp3* ^60^), meiotic transcriptional program, such as *Brdt* ^8^), and meiotic cell cycle, such as *Meioc* ^61^) (Figure 4G). Taken together, these findings indicate that the NFYA-SMARCA5-BPTF axis activates the mitotic transcriptional program required for spermatogonial differentiation and meiotic entry, while SMARCA5 can also act independently of NFYA and BPTF to repress genes that should remain silent during this developmental transition.

### NFYA and the NURF complex co-occupy poised meiotic gene promoters in spermatogonia

We found that NFYA interacts with BPTF and SMARCA5, i.e., the NURF complex (Figure 1A). In fact, SMARCA5 occupancy at poised meiotic promoters, but not at mitotic promoters, was markedly reduced in NFYA-deficient SpG (Figure 1F), indicating that NFYA recruits NURF complex specifically to poised meiotic gene promoters. We thus sought to directly determine whether NFYA and NURF complex cooperatively bind to the promoters of poised meiotic genes in SpG. Remarkably, in SpG, we detected strong NFYA, SMARCA5, and BPTF occupancies around the transcription start sites (TSSs) of poised genes. In contrast, little to no NFYA, SMARCA5, or BPTF signal was detected around the TSSs of non-poised genes (Figures 5A and S8). Indeed, we found moderate-to-strong correlation between chromatin accessibility and the occupancies of NFYA, SMARCA5, or BPTF, such that higher chromatin accessibility was associated with greater occupancy of each factor (Figure 5B; Spearman’s ρ = 0.49 – 0.71). Likewise, the occupancies of NFYA, SMARCA5, and BPTF were strongly correlated (Figure 5B; Spearman’s ρ = 0.62 – 0.91). Representative examples of poised and non-poised genes corroborated these findings: promoters of poised genes exhibited accessible chromatin and were occupied by NFYA, SMARCA5, and BPTF, whereas promoters of non-poised genes lacked both accessible chromatin and NFYA, SMARCA5, BPTF occupancy (Figure 5C).

**Figure 5.**
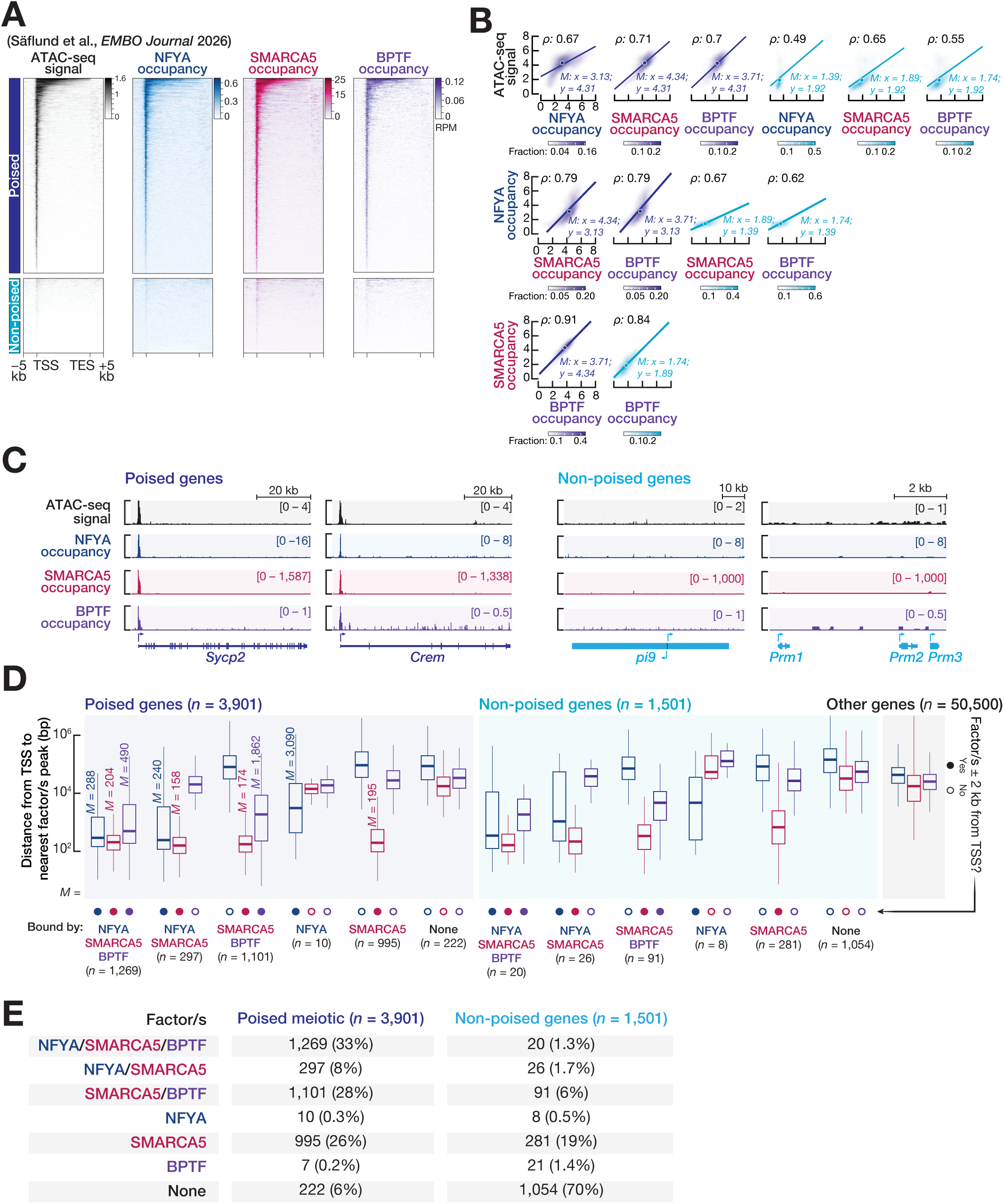
NFYA and the NURF complex co-occupy poised meiotic gene promoters in spermatogonia. See also Figure S8. **(A)** Heatmaps show the reads per million (RPM)-normalized ATAC-seq signal, together with NFYA, SMARCA5, and BPTF occupancy, in the −5 kb to +5 kb regions flanking the transcription start sites (TSSs) and transcription end sites (TESs) of poised and non-poised genes in spermatogonia (SpG). One of the three biological replicates is shown; the remaining replicates are presented in Extended Data Figure 8. **(B)** Scatter plots show the Spearman’s correlations (ρ) between chromatin accessibility and NFYA, SMARCA5, or BPTF occupancy, as well as among the occupancies of NFYA, SMARCA5, and BPTF at the promoters of poised genes in SpG. Each data point represents the average signal from three biological replicates. **(C)** Integrative Genomics Viewer show RPM-normalized ATAC-seq signal, together with RPM-normalized NFYA, SMARCA5, and BPTF occupancy, at the gene boundaries of exemplified poised (*Sycp2* and *Crem*) and non-poised (*pi9*, *Prm1*, *Prm2*, and *Prm3*) genes in SpG. **(D)** Box plots show the distance (average of three biological replicates) from the TSS to the nearest NFYA, SMARCA5, or BPTF peak for poised genes (*n* = 3,901), non-poised genes (*n* = 1,501), and other (*n* = 50,500) genes excluding poised, non-poised, and mitotic genes in SpG. Genes are classified into four categories based on NFYA, SMARCA5, and BPTF occupancy: genes co-bound by all three factors, genes co-bound by SMARCA5 and BPTF, genes bound by SMARCA5 alone, and genes not bound by any of these factors. A Gene was considered as factor-bound if a significant peak was detected within ±2 kb of its TSS in at least two biological replicates. Horizontal lines indicate the median. Whiskers show the minimum and maximum values. Boxes represent the interquartile range (IQR). **(E)** Table summarizes the numbers and percentages of poised and non-poised genes with promoters co-occupied by all three factors, co-occupied by SMARCA5 and BPTF, occupied by SMARCA5 alone, or not occupied by any of these factors.

We next determined the number of poised genes bound by NFYA, SMARCA5, and BPTF in SpG. Notably, the promoters of approximately 33% of poised genes were bound by all three factors (Figures 5D, E and Table S5; 1,269 of 3,901 genes with median distances from TSS to the nearest NFYA, SMARCA5, BPTF peaks of 288 bp, 204 bp, 490 bp, respectively). In contrast, the promoters of fewer than 2% of non-poised genes were bound by all three factors (Figures 5D, E; 20 of 1,501 genes). Furthermore, nearly 28% of poised gene promoters were occupied by SMARCA5 and BPTF in the absence of NFYA binding (Figures 5D, E), suggesting that recruitment of the NURF complex to these promoters is likely mediated by another factor. Given that NFYA is required to maintain chromatin accessibility at poised meiotic promoters prior to meiotic entry^22^, the co-occupancy of NFYA and its interacting partner, the NURF complex, at these promoters in SpG suggests that NFYA and NURF cooperate to regulate chromatin accessibility.

### The NFYA-NURF axis is required to maintain accessible chromatin at poised meiotic promoters

To directly test whether the NURF complex, in collaboration with NFYA, regulates permissive chromatin at poised meiotic promoters, we examined the change in chromatin accessibility at promoters of poised in SMARCA5-deficient SpG compared to *Cre/+* SpG. We analyzed the scATAC-seq dataset of our sc-multiomics experiment. After filtering doublets, we clustered 64,361 cells (average TSS enrichment score per cell = 10.15; Supplemental Table S6) from 78,860 individual cells whose identities were resolved in scRNA-seq (Figure 3D), using a genome-wide 500-bp binarized tile matrix with ArchR^62^ (Figure 6A). The resulting UMAP recapitulated complete germ cell developmental trajectory (Figure 6A), as exemplified by distinct chromatin accessibility around the TSSs of several representative cell-type-specific marker genes across major cell populations (Figure 6B).

**Figure 6.**
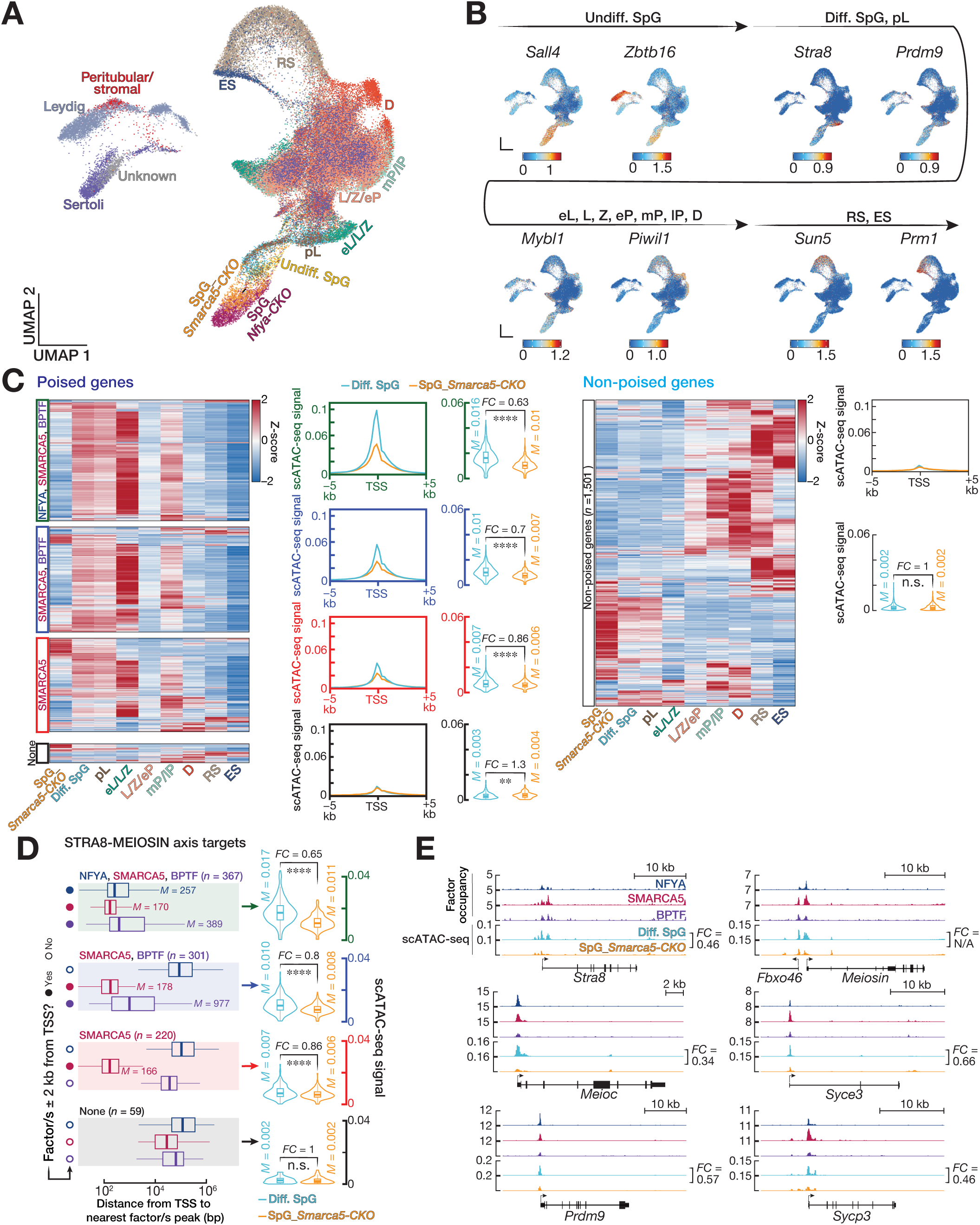
the NFYA-NURF axis regulates permissive chromatin at poised meiotic promoters in pre-meiotic spermatogonia. **(A)** Dimension reduction analysis by uniform manifold approximation and projection (UMAP) on 64,361 individual cells captured by combined scATAC- and scRNA-seq datasets from 8-weeks old *Cre/+*, *Smarca5-CKO* and *Nfya-CKO* testes identified 14 cell populations. **(B)** Per-cell gene ATAC-seq signal patterns of exemplified marker genes are visualized in UMAP. Direction of germ cell developmental trajectory is indicated by black arrows. **(C)** Heatmaps show relative ATAC-seq signal (Z-score) in the −2 kb to +2 kb regions flanking the transcription start sites (TSSs) of poised (left) and non-poised (right) genes in different cell population defined by scRNA-seq. Poised genes are divided into four categories: NFYA-NURF axis-bound (*n* = 1,224), the NURF complex-bound (*n*□=□1,073), SMARCA5-alone-bound (*n* = 962), and not bound by any of the three factors (*n* = 222). Metagene plots of ATAC-seq signal in the −5 kb to +5 kb regions flanking the TSSs of poised (left) and non-poised (right) genes. Violin plots show the ATAC-seq signal in the −2 kb to +2 kb regions flanking the TSSs of four poised gene categories (left) and non-poised genes (right) in SMARCA5-deficient SpG compared to diff. SpG of *Cre/+.* FC, Fold Change. \*\*\*\**p* < 0.0001, \*\**p* < 0.01, n.s., not significant. Significance was measured by two-sided Wilcoxon matched-pairs signed-ranked sum test. **(D)** Box plots (left) show the distance (average of three biological replicates) from the TSS to the nearest NFYA, SMARCA5, or BPTF peak for STRA8-MEIOSIN axis target genes in SpG. Genes are classified into four categories based on NFYA, SMARCA5, and BPTF occupancy: NFYA-NURF axis-bound (*n* = 367), the NURF complex-bound (*n*□=□301), SMARCA5-alone-bound (*n*□=□220), and not bound by any of the three factors (*n* = 59). Whiskers show the minimum and maximum values. Boxes represent the interquartile range (IQR). Violin plots show the ATAC-seq signal in the −2 kb to +2 kb regions flanking the TSSs of four STRA8-MEIOSIN target gene categories in SMARCA5-deficient SpG compared to diff. SpG of *Cre/+.* FC, Fold Change. \*\*\*\**p* < 0.0001, n.s., not significant. Significance was measured by two-sided Wilcoxon matched-pairs signed-ranked sum test. **(E)** Integrative Genomics Viewer (IGV) tracks (left) show the reads per million (RPM)-normalized NFYA, SMARCA5, BPTF occupancy, together with scATAC-seq signals from SMARCA5-deficient SpG and diff SpG of *Cre/+*, at gene boundaries of exemplified genes controlled by STRA8-MEIOSIN regulatory axis. FC, Fold Change, denotes the change in scATAC-seq signal between SMARCA5-deficient SpG and diff. SpG of *Cre/+*.

Poised meiotic promoters bound by the NFYA-NURF axis, the NURF complex alone, and SMARCA5 alone displayed reduced chromatin accessibility in SMARCA5-deficient SpG when compared to *Cre/+* SpG. By contrast, poised promoters not occupied by any of these factors, as well as non-poised promoters remained largely unchanged between SpG from *Smarca5-CKO* and *Cre/+* mice (Figure 6C). Consistent with a cooperative role of NFYA and the NURF complex, the reduction in chromatin accessibility at poised meiotic promoters was greater at the NFYA-NURF axis-occupied promoters that at promoters bound by either the NURF complex alone or SMARCA5 alone (Figure 6C; NFYA-NURF axis-bound promoters, *Smarca5-CKO* ÷ *Cre/+* = 0.63; the NURF complex-bound promoters, *Smarca5-CKO* ÷ *Cre/+* = 0.7; SMARCA5-alone-bound promoters, *Smarca5-CKO* ÷ *Cre/+* = 0.86; two-sided wilcoxon matched-pairs signed-ranked sum test).

NFYA regulates chromatin accessibility at the promoters of *Stra8* and *Meiosin* and at genes controlled by STRA8-MEIOSIN transcriptional axis ^22^. We therefore investigated whether NFYA and the NURF complex collaboratively regulate permissive chromatin at the promoters of STRA8-MEIOSIN target genes. Notably, chromatin accessibility at *Stra8* and *Meiosin* promoters, both of which are bound by the NFYA-NURF axis, was reduced in SMARCA5-deficient SpG (Figure 6D, E). Of the 1,044 STRA8-MEIOSIN target genes, 367, 301, and 220 promoters were bound by the NFYA-NURF axis, the NURF complex alone, and SMARCA5 alone, respectively (Figure 6D). Although chromatin accessibility was reduced at all three promoters categories in SMARCA5-deficient SpG, the reduction was most pronounced at NFYA-NURF axis-bound promoters (Figure 6D; NFYA-NURF axis-bound promoters, *Smarca5-CKO* ÷ *Cre/+* = 0.65; the NURF complex-bound promoters, *Smarca5-CKO* ÷ *Cre/+* = 0.8; SMARCA5-alone-bound promoters, *Smarca5-CKO* ÷ *Cre/+* = 0.86; two-sided wilcoxon matched-pairs signed-ranked sum test). These findings support a cooperative role of NFYA and the NURF complex in maintaining chromatin accessibility at poised meiotic promoters. Notably, genes, whose promoters were bound by the NFYA-NURF axis and exhibited reduced chromatin accessibility in SMARCA5-deficient SpG, included key regulators of meiotic recombination, such as *Meioc*, *Prdm9*, *Syce3*, and *Sycp3* (Figure 6E). We conclude that the NFYA-NURF axis is essential for the maintenance of chromatin accessibility at meiotic promoters, including those activated by the STRA8-MEIOSIN axis, in pre-meiotic SpG.

### Combinatorial occupancy by NFYA, NURF, SMARCA5, ZBTB16 and SALL4 predicts the temporal activation of poised meiotic genes

We found that NFYA and the NURF complex co-occupy one-third of poised promoters, whereas the remaining two-thirds are occupied in approximately equal proportions by either the NURF complex alone or SMARCA5 alone (Figure 5). The ZBTB16-SALL4-SOX3 regulatory axis, enriched in undiff. SpG ^45,63^, is recently shown to help establish poising at a large fraction of meiotic gene promoters in undiff. SpG^23^. This prompted us to investigate what distinguishes poised meiotic promoters bound by NFYA-NURF axis from those occupied by either the NURF complex alone or SMARCA5 alone. We re-analyzed published ZBTB16 and SALL4 ChIP-seq datasets from 7 dpp mouse testes^23^ to examine the functional interplay between the ZBTB16-SALL4 regulatory axis and the NFYA-NURF axis, the NURF complex, and SMARCA5 prior to meiotic initiation. Genome-wide occupancy and peak analyses revealed that promoters not bound by the NFYA-NURF axis, the NURF complex, and SMARCA5 alone, were likewise devoid of ZBTB16 and SALL4 binding (Figure 7A), suggesting that these factors converge on the regulation of a common set of genes. Nevertheless, among poised promoters, those lacking the occupancy of NFYA-NURF remained bound by ZBTB16 and SALL4 (Figure 7B, left; Mann–Whitney–Wilcoxon *U* test), consistent with the widespread occupancy of ZBTB16-SALL4-SOX3 regulatory axis at poised promoters before the onset of meiosis^23^. Notably, however, promoters bound by SMARCA5 alone exhibited lower H3K27Ac signals but higher levels of repressive mark H2AK119ub than promoters occupied by the NFYA-NURF axis (Figure 7B, left; Mann–Whitney–Wilcoxon *U* test), suggesting that, despite their poised state, such genes are perhaps actively repressed to prevent premature transcription at inappropriate developmental stage after meiotic entry. Consistent with this idea, following meiotic entry, the expression of poised genes bound exclusively by SMARCA5 and marked by higher H2AK119ub signals peaked later than that of genes occupied by the NFYA-NURF axis (Figure 7C). Furthermore, GO analysis showed that genes bound by SMARCA5 alone were enriched for processes related for spermiogenesis, including motile cilium assembly, cilium-dependent cell motility, sperm motility, spermatid development. In contrast, genes bound by the NFYA-NURF axis were enriched for processes required for meiotic entry and progression, such as centriole replication, positive regulation of DSB, positive regulation of DNA recombination (Figure 7D). Together, these findings suggest that multiple factors form a regulatory network that regulate poised state of meiotic promoters in pre-meiotic germ cells, and that the precise temporal expressions of poised genes depend on the combinatorial binding of NFYA, the NURF complex, SMARCA5, and ZBTB16-SALL4 axis (Figure 7E).

**Figure 7.**
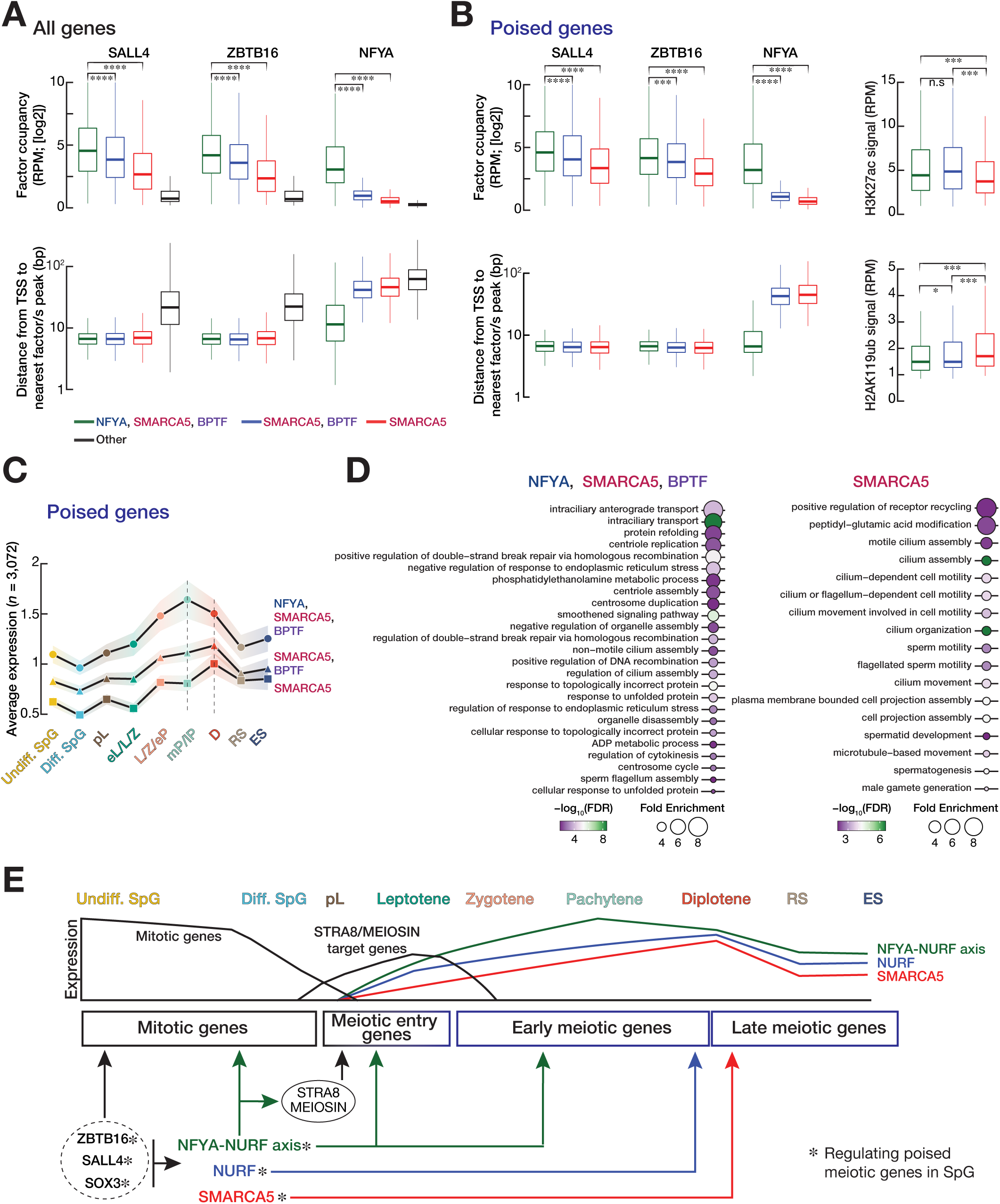
Combinatorial occupancy of the NFYA-NURF axis, the NURF complex, SMARCA5, ZBTB16, and SALL4 at poised meiotic gene promoters predicts their timely activation. **(A)** Box plots (top) show the occupancies of SALL4, ZBTB16 and NFYA (average of two or three replicates) around the transcription start sites (TSSs) of genes bound by the NFYA-NURF axis (*n*□=□4,700), the NURF complex (*n*□=□5,044), SMARCA5 alone (*n*□=□6,548) and genes unbound by any of these factors (*n* = 39,208) in prepubertal testis. Publicly available ZBTB16 and SALL4 ChIP-seq datasets from 7 dpp mouse testes^23^ were re-analyzed. Box plots (bottom) shows the distance (average of two or three biological replicates) from the TSS to the nearest SALL4, ZBTB16, or NFYA peak for genes bound by the NFYA-NURF axis, the NURF complex, SMARCA5 alone and genes unbound by any of these factors. \*\*\*\**p* < 0.0001, n.s. not significant. Statistical significance is measured by Mann–Whitney–Wilcoxon *U* test. Horizontal lines indicate the median. Whiskers show the minimum and maximum values. Boxes represent the interquartile range (IQR). **(B)** Box plots (top left) show the occupancies of SALL4, ZBTB16 and NFYA (average of two or three replicates) around the TSSs of poised genes bound by the NFYA-NURF axis (*n*□=□1,224), the NURF complex (*n*□=□1,073) or SMARCA5 alone (*n*□=□962) in prepubertal testis. Box plots (bottom left) shows the distance (average of two or three biological replicates) from the TSS to the nearest SALL4, ZBTB16, or NFYA peak for poised genes bound by the NFYA-NURF axis, the NURF complex, SMARCA5 alone. Box plots (top right and bottom right) show the RPM-normalized H3K27ac and H2AK119ub occupancy around the TSSs of poised genes bound by the NFYA-NURF axis, the NURF complex or SMARCA5 alone in diff. SpG. \**p* < 0.05; \*\*\**p* < 0.001. Statistical significance is measured by Mann–Whitney–Wilcoxon *U* test. Horizontal lines indicate the median. Whiskers show the minimum and maximum values. Boxes represent the IQR. **(C)** Line plot of average expression of poised genes bound by the NFYA-NURF axis (*n*□=□1,159), the NURF complex (*n*□=□1,019), or SMARCA5 alone (*n* = 894) in nine different cell populations defined by scRNA-seq. **(D)** Geno Ontology (GO) analysis for poised genes bound by the NFYA-NURF complex (left) and bound by SMARCA5 alone (right). Top gene categories with ≥2-fold enrichment and FDR < 0.01 are shown. **(E)** The model illustrates the regulatory network of factors that cooperatively control the temporal expression of genes during spermatogenesis. The model is constructed based on this and previous studies^4,5,7–13,22,23,67,68^. The model emphasizes the sequential and coordinated functions of key regulatory factors, i.e., SOX3, SALL4, ZBTB16, NFYA, SMARCA5, STRA8 and MEIOSIN throughout the developmental progression of male germ cells.

## Discussion

The establishment of accessible chromatin at developmental genes before their transcription is a recurring feature of developmental gene regulation^64^. During spermatogenesis, before the onset of meiosis, many meiotic genes acquire accessible chromatin accompanied by accumulation of paused RNA polymerase II (Pol II), thereby generating a poised regulatory state that permits timely activation of meiotic genes upon meiotic entry^8,21–23^. Although this poised state is recognized as a hallmark of male germ cell development, the molecular mechanisms responsible for regulating this permissive chromatin landscape at poised meiotic promoters is poorly understood. Here, we identify the NURF chromatin-remodeling complex, comprised of the ISWI ATPase SMARCA5 and its accessory subunit BPTF, as a key regulator of this process and demonstrate its functional dependency on NFYA. Our findings support a model in which NFYA is required for the NURF complex occupancy at poised meiotic promoters prior to meiotic entry, thereby facilitating permissive chromatin environment that primes meiotic genes for subsequent activation during meiosis (Figure 7E).

Pioneer transcription factors are distinguished by their ability to engage closed or partially closed chromatin and initiate local chromatin remodeling; in diverse developmental systems, pioneer factors collaborate with ATP-dependent chromatin remodeling complexes to establish the competency of regulatory elements before developmental genes become activated^26–29^. Our findings place NFYA within this conceptual framework. We show that NFYA interacts with SMARCA5, that SMARCA5 occupancy at poised meiotic promoters is largely dependent on NFYA, and that NFYA and the NURF complex co-occupy the substantial fraction of poised meiotic gene promoters in SpG. In addition to the enrichment of NFYA-binding motif within the promoters of poised meiotic genes^22^, NFYA forms the NF-Y complex together with NFYB and NFYC^65^. The NF-Y complex induces nearly 80° bend in DNA, thereby facilitating the binding of additional regulatory factors^66^, suggesting that NFYA may also recruit other factors by acting as a molecular scaffold. Furthermore, NFYA has been shown to bind inaccessible chromatin in other cellular contexts^24^. Together, our work and previous studies support a model in which NFYA functions as a pioneer factor in male germ cells.

In addition to the establishment of chromatin accessibility at poised developmental gene promoters, maintenance of permissive chromatin is an equally important process that ensures timely activation of developmental genes^26–29^. Although the establishment of accessible chromatin has received considerable attention in developmental systems, less is known about how this permissive state is maintained during extended developmental intervals before genes become transcriptionally active. Our findings together with previous studies support a stepwise model for the regulation of meiotic genes. ZBTB16-SALL4-SOX3 regulatory axis first establishes promoter poising in undiff. Spg^23^, the NFYA-NURF regulatory axis maintains accessible chromatin in diff. SpG, and the STRA8-MEIOSIN and AMYB-TCFL5 regulatory axes subsequently activate the meiotic transcriptional program. Therefore, the developmental competence of poised meiotic promoters is actively preserved by continuous ATP-dependent chromatin remodeling throughout spermatogonial differentiation prior to meiotic entry.

Approximately two-thirds of poised meiotic promoters are not occupied by the NFYA-NURF axis. Among these promoters, nearly half are bound by the NURF complex in the absence of NFYA, whereas the remaining half exhibit SMARCA5 occupancy alone (Figure 5). Our re-analyses of ZBTB16 and SALL4 ChIP-seq datasets revealed occupancy of both factors at these NFYA-independent promoters, suggesting that poising of meiotic promoters is not controlled by a single regulatory axis but rather involves multiple factors, together with additional yet-to-be-identified factor/s, that converge on a common chromatin remodeling machinery. Furthermore, we found that poised genes with promoters occupied exclusively by SMARCA5 are enriched for gene categories related to spermiogenesis and are expressed later than those occupied by the NFYA-NURF axis during meiosis. Consistent with this delayed activation, promoters bound by SMARCA5 alone display a more repressive chromatin environment, characterized by increased H2K119ub occupancy and reduced H3K27Ac (Figure 7B). Together, these observations suggest that combinatorial occupancy by the NFYA-NURF axis, the NURF complex, SMARCA5, and the ZBTB16-SALL4 axis not only regulates promoter poising but also perhaps encode information that predicts the temporal activation of meiotic genes during meiosis.

Finally, ISWI chromatin remodelers can mediate gene repression through their association with alternative accessory proteins^31,49^. Consistently, we observed increased expression of many genes associated with biological processes related to non-germ cell differentiation and macromolecule metabolism in SMARCA5-deficient SpG (Figures 4A, B), suggesting that SMARCA5 represses their transcription in male germ cells. Notably, such gene promoters bound by SMARCA5 alone exhibit higher repressive chromatin environment compared to those occupied by the NFYA-NURF axis (Figure 4D). A recent study showed that SMARCA5 restricts permissive chromatin at numerous promoters required for meiotic progression^67^, supporting a broader role for SMARCA5 beyond transcriptional activation during spermatogenesis. It is possible that distinct SMARCA5-accessory protein assemblies may either mediate transcriptional activation or repression at different classes of promoters. Elucidating the roles of other potential accessory proteins of SMARCA5 in spermatogonial differentiation would be an interesting avenue for future studies.

In summary, our work reveals the developmental competence at poised meiotic promoters is actively maintained by the NFYA-NURF regulatory axis in spermatogonia. Findings in our study extend the concept of promoter poising beyond its establishment and highlight the importance of continuous ATP-dependent chromatin remodeling activity to preserve meiotic poising before meiotic genes become activated. We propose that active maintenance of permissive chromatin is a unifying feature of developmental systems that ensures timely execution of cell-fate transitions.

### Limitations of the study

Our study identifies the NFYA-NURF axis as a key regulator of chromatin accessibility at poised meiotic genes in pre-meiotic spermatogonia. However, we acknowledge some limitations of our studies. Although we demonstrate that SMARCA5 occupancy at poised meiotic promoters depends on NFYA, whether the interaction between NFYA and SMARCA5 is direct remains to be determined. It is unclear whether NFYA recruits SMARCA5 through a direct physical interaction or indirectly via one or more intermediary factors. Although our work supports a role for the NURF complex – comprised of SMARCA5 and BPTF – in maintaining chromatin accessibility, we did not examine the consequences of loss of BPTF in germ cells. Future studies are warranted to examine the extent to which BPTF contributes to permissive chromatin state at poised meiotic promoters.

## Resource availability

### Lead contact

Request for further information and data availability will be processed by the lead contact Deniz M. Özata.

### Material availability

Sequencing data are available on the Gene Expression Omnibus (GEO) with the accession number, GSE349954. The raw MS data are available on the jPOST Repository with the accession number, PXD083858. Microscopy and western blotting images are deposited on BioStudies with the accession number, S-BIAD4123.

### Data and code availability

All scripts used to analyze sequencing data can be found at the Ozata lab GitHub: https://github.com/Ozatalab. All publicly available datasets were downloaded from the GEO:

- CUT&Tag for H2AK119ub from differentiating spermatogonia^51^: GSE221942.
- H3K27ac ChIP-seq from type B spermatogonia^50^: GSE132446.
- SALL4 ChIP-seq from seven days old mice testis^23^: GSE202819.
- ZBTB16 ChIP-seq from seven days old mice testis ^23^: GSE202819.
- ATAC-seq from FACS-sorted spermatogonia^22^: GSE295991.
- CUT&RUN for NFYA from FACS-sorted spermatogonia^22^: GSE295991.
- scRNA-seq and scATAC-seq from the testicular nuclei of *Cre/+* and *Nfya-CKO* mice22: GSE295991.

## Acknowledgements

We thank the personnel of Stockholm University animal facility, with particular gratitude to S. Olsson, S. Oerther, and R. Askar for their excellence in mouse colony management; C. Molenaar for his expert help with Zeiss Axio Observer 7 inverted widefield microscope; National Genomics Infrastructure—funded by Science for Life Laboratory, the Knut and Alice Wallenberg Foundation and the Swedish Research Council.

## Authors’ contributions

D.M.Ö. and Y.T.X. conceived and designed the experiments. A.E., M.S., and M.M.A. performed the experiments. M.A. and M.S. analyzed the sequencing data. A.E., M.S., M.M.A., and D.M.Ö. characterized the *Smarca5-CKO* mice phenotype. A.E., M.S., M.A., AK.I.Ö.F., Y.T.X., and D.M.Ö. wrote the manuscript. All authors discussed and approved the manuscript.

## Supplemental information titles and legends

**Document S1.** Supplemental Figures S1–S8

**Table S1:** Mass spectrometry results.

**Table S2:** Statistics of scRNA-seq experiment.

**Table S3:** Differential gene expression analysis of scRNA-seq.

**Table S4:** Gene ontology analyses for deregulated genes in SMARCA5-deficient SpG.

**Table S5:** The distances from the TSSs of poised genes to the nearest NFYA, SMARCA5, and BPTF peaks.

**Table S6:** Statistics of scATAC-seq experiment.

## STAR Methods

### Key resources table

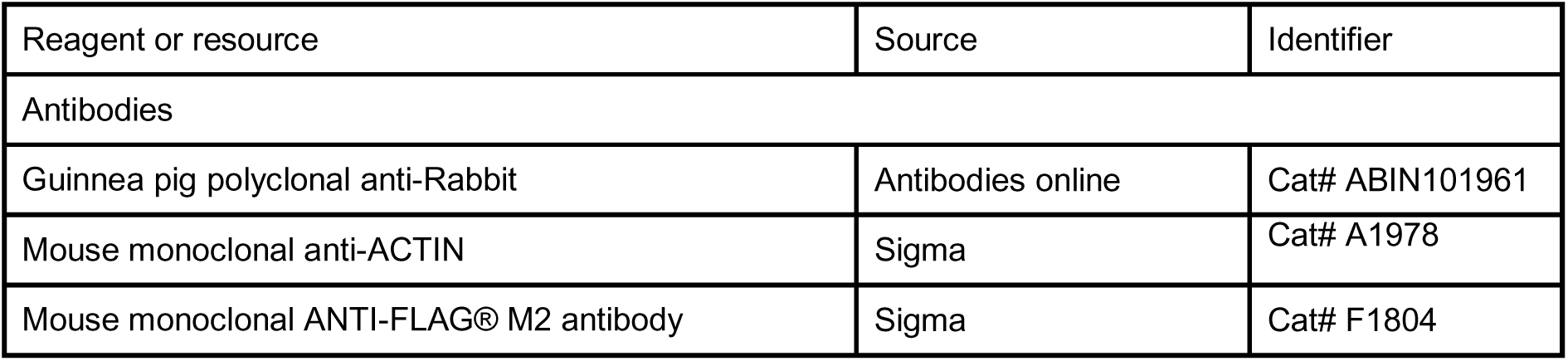
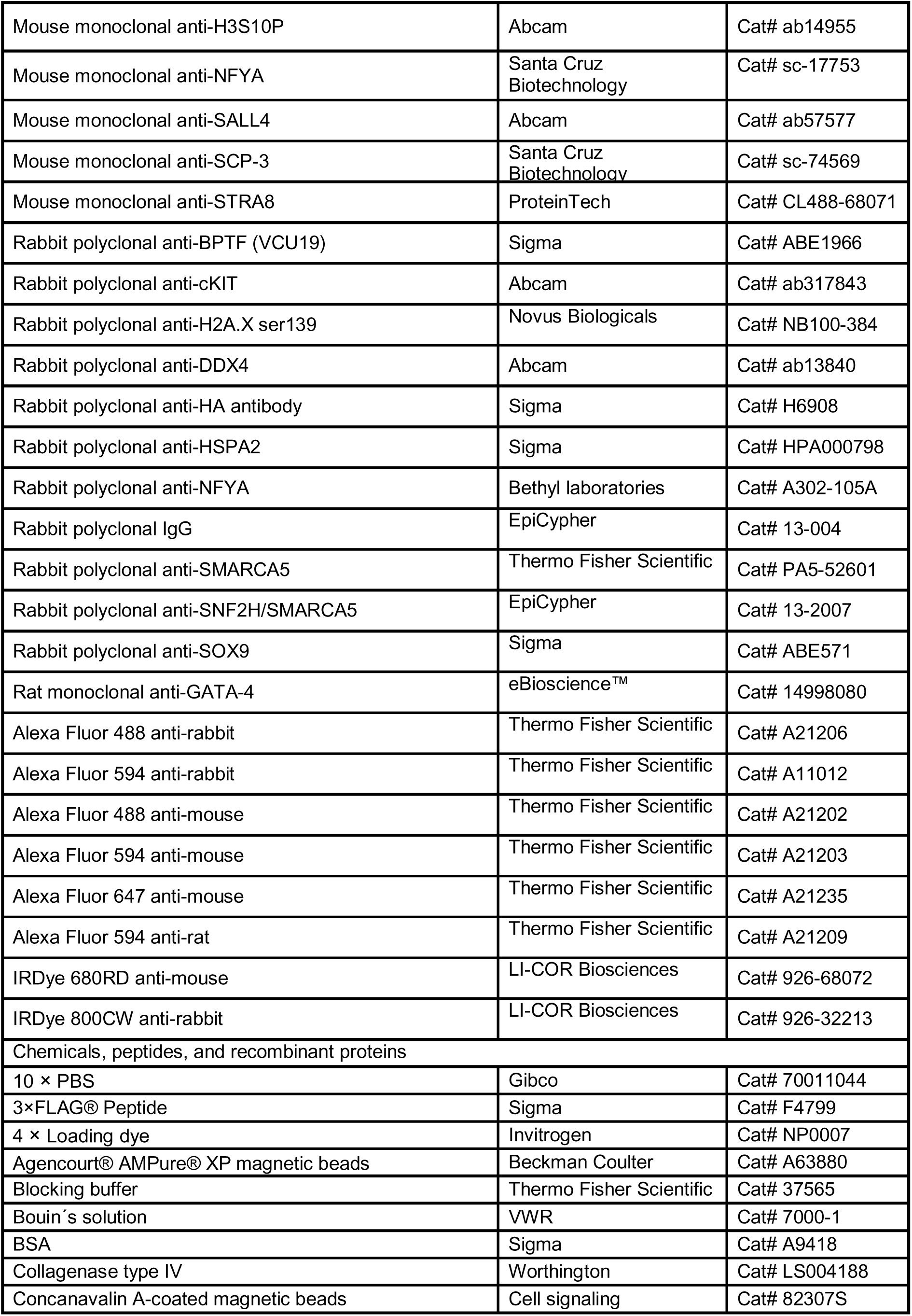
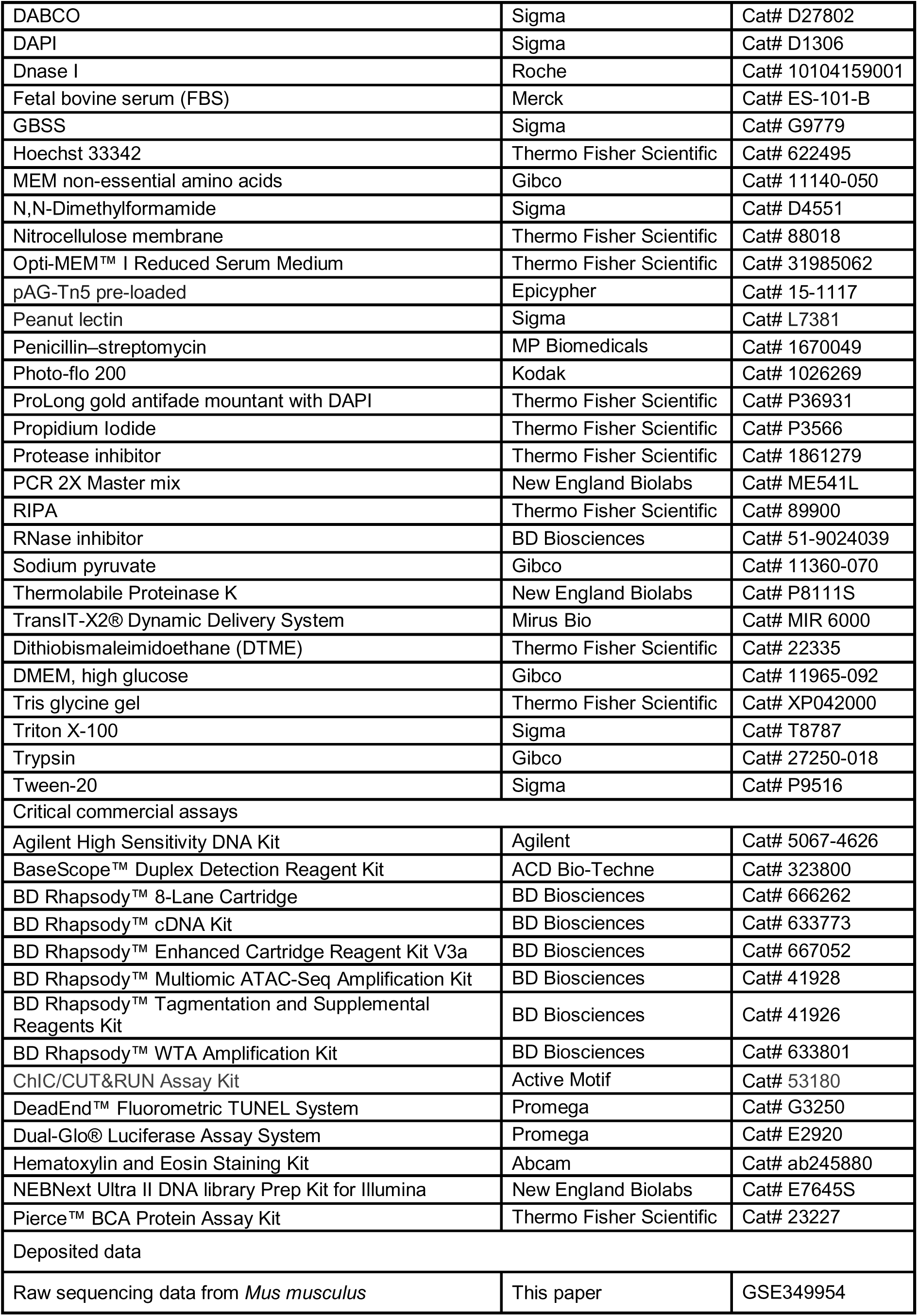
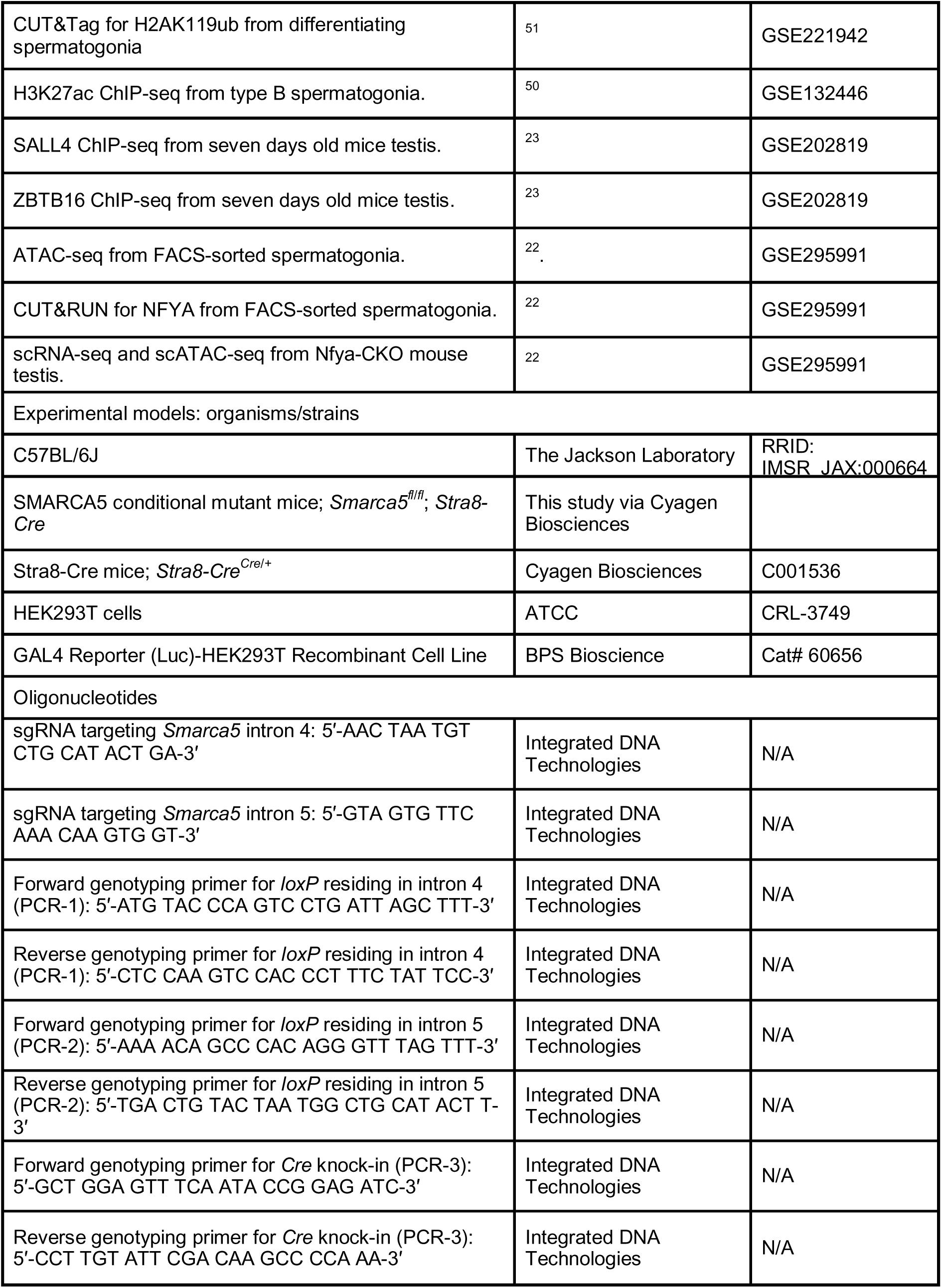
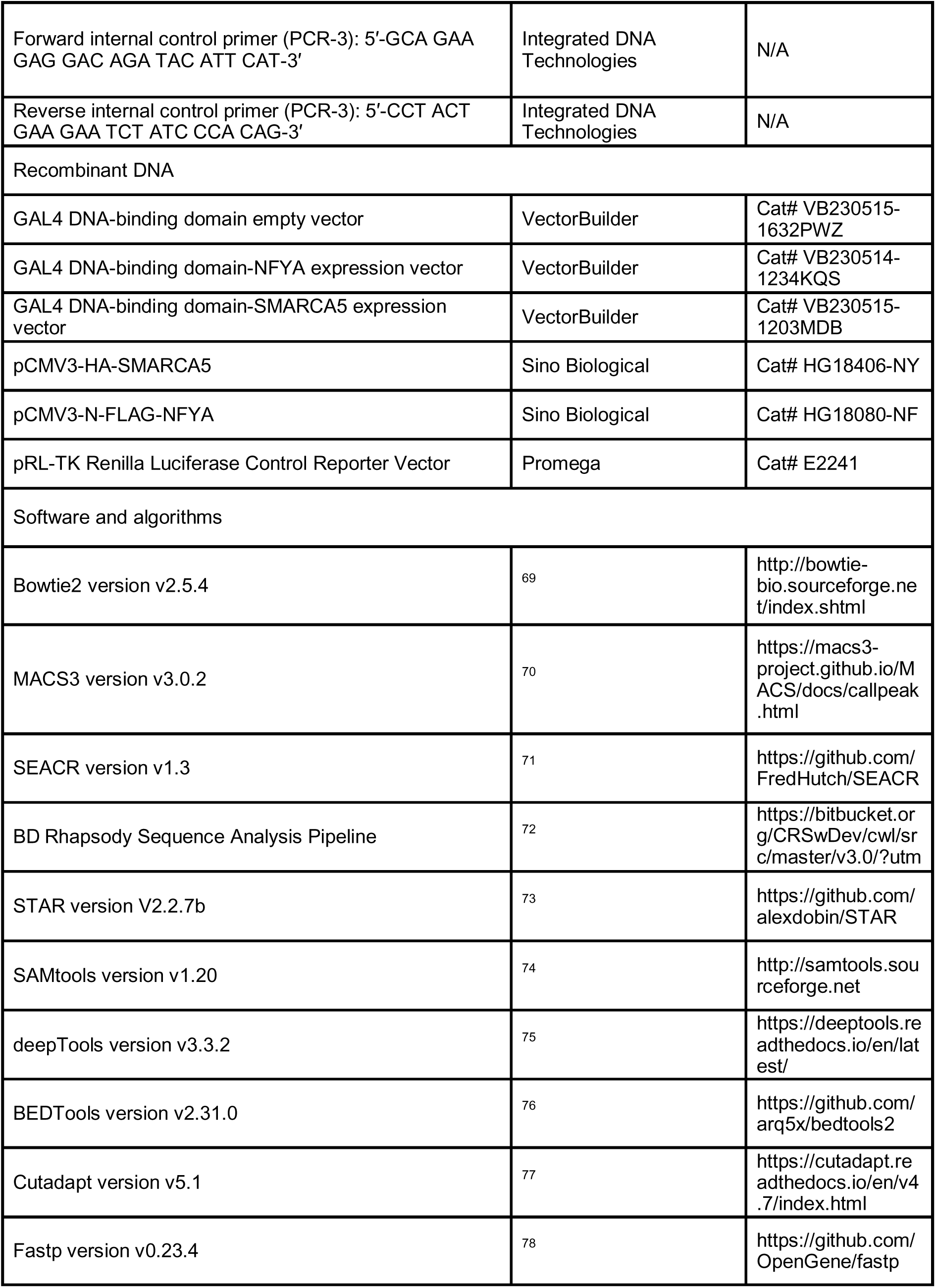
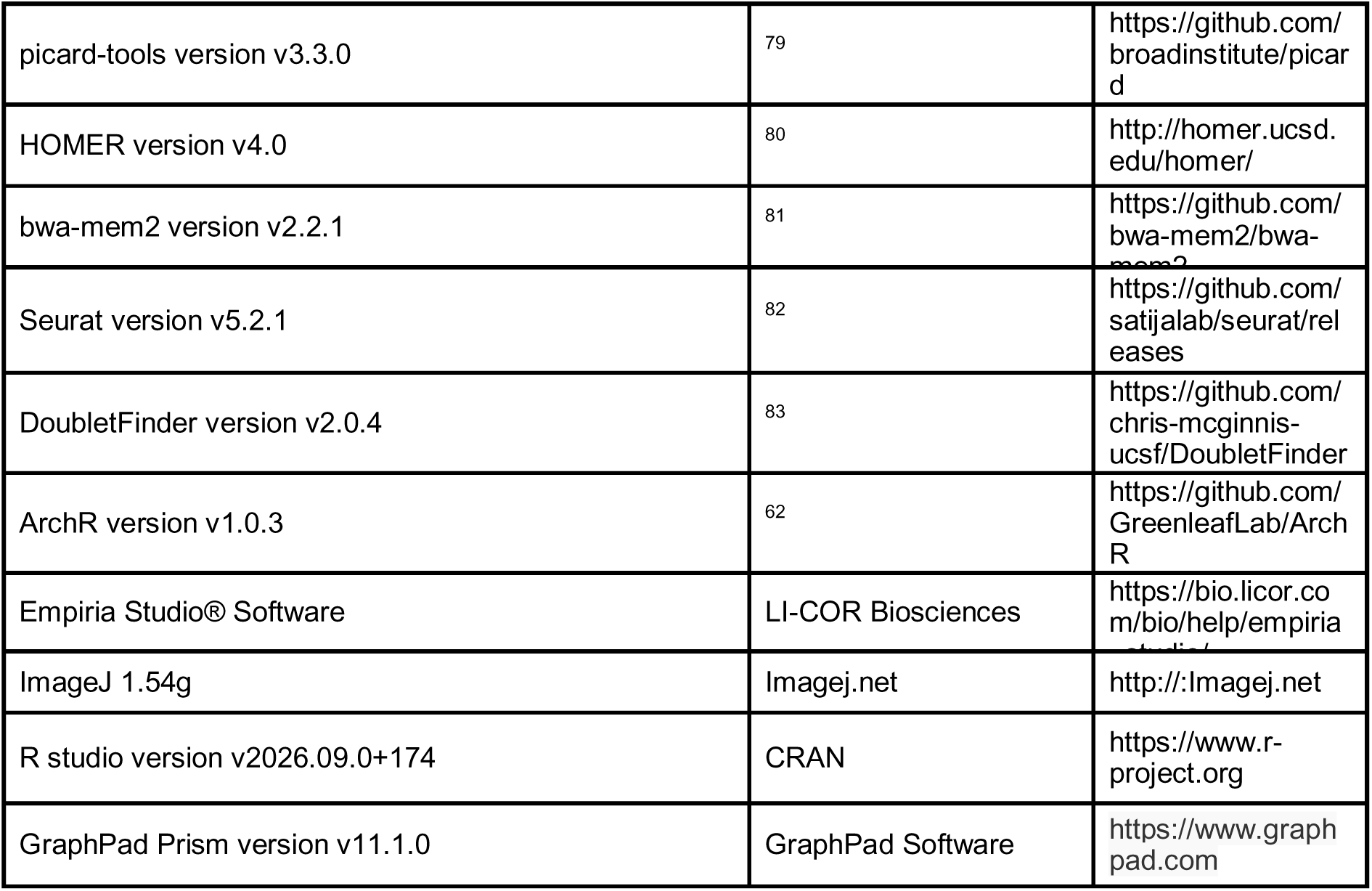

### Experimental models and study participants

#### Mice

Animal experiments were conducted in accordance with the guidelines established by Stockholm University and approved by the Regional Animal Experimentation Ethics Committee of the Swedish Board of Agriculture with the animal ethical permit numbers 8246-2021 and 5.8.18-3736/2026. C57BL/6J mice (RRID: IMSR_JAX:000664) were used as wild-type control. *Smarca5* mutant mice were generated by conditional deletion of exon 5 of *Smarca5* using the Stra8-Cre-loxP system (Cyagen Biosciences, Jiangsu, China). Stra8-P2A-ZsGreen1-T2A-Cre mice were obtained from Cyagen (Cyagen Biosciences, C001536). The 5′ and 3′ loxP sites were inserted into introns 4 5, respectively. Briefly, two guide RNAs targeting sequences in intron 4 (5′-AAC TAA TGT CTG CAT ACT GA-AGG-3′) and intron 5 (5′-GTA GTG TTC AAA CAA GTG GT-TGG-3′), together with a donor vector containing loxP sites and Cas9 protein, were co-injected into the cytoplasm of pronuclear-stage embryos. The injected embryos were cultured overnight in KSOM medium, and the resulting two-cell-stage embryos were subsequently transferred into the uterus of pseudopregnant ICR females. F0 founder mice were then backcrossed with C57BL/6J mice to identify animals exhibiting germline transmission in the F1 generation.

#### Cell lines

HEK293T cells (ATCC, CRL-3749) were maintained in high-glucose Dulbecco’s modified Eagle’s medium (DMEM; Gibco, 11965-092) supplemented with 10% fetal bovine serum (FBS; Merck, ES-101-B), 1% penicillin–streptomycin (MP Biomedicals, 1670049), 1% MEM non-essential amino acids (Gibco, 11140-050), and 1% sodium pyruvate (Gibco, 11360-070) at 37°C in a humidified atmosphere containing 5% CO_₂_.

## Method details

### Mass spectrometry

#### Immunoprecipitation

A frozen testis tissue was minced into small pieces and thoroughly homogenized in RIPA lysis buffer (Thermo Fisher Scientific, 89900, 25 mM Tris-HCl, pH 7.6, 150 mM NaCl, 1% (v/v) NP-40, 1% (w/v) sodium deoxycholate, and 0.1% (w/v) SDS) supplemented with 1% (v/v) protease inhibitor (Thermo Fisher Scientific, 1861279) and 500 mM NaCl using 20–30 strokes of pestle B. The samples were then centrifuged at 21,000g in 4°C for 30 min. The supernatant was collected into a new tube, and two volumes of complete RIPA buffer was added to reduce the salt concentration. The protein lysate was then precleared with 50 μl Dynabeads Protein G (Thermo Fisher Scientific, 10003D) at 4°C for 1h. After preclearing, the lysate was divided into two tubes: 2 μg anti-NFYA antibody (Bethyl, A302-105A) was added to one tube, and rabbit anti-IgG antibody (Abcam, ab172730) was added to the other. The lysates were incubated with the primary antibodies overnight at 4°C. Next day, 50 μl Dynabeads Protein G were added to each tube and incubated at 4°C for 1 h. The antibody-bead complexes were washed three times with complete RIPA buffer before elution with 0.1M acidic glycine pH 2.0 or with complete RIPA and Loading dye (Invitrogen, NP0007) containing 0.2 M dithiothreitol (DTT). Western blotting was performed as described below, or the samples were processed for LC-MS/MS.

#### LC-MS/MS analysis

LC-MS/MS digestion and analysis were performed by the University of Massachusetts Proteomics Core as previously described^84^. Briefly, eluates from NFYA immunoprecipitation were denatured in loading dye (Invitrogen, NP0007) containing 0.2M dithiothreitol (DTT) at 95°C for 5 min and resolved through a 4–20% Tris-Glycine gradient SDS gel (ThermoFisher Scientific, XP04200) for 10 min at 100 V. The entire area above bromophenol blue and xylene cyanol was excised. The gel pieces were then reduced with DTT and alkylation and were subjected to trypsin digestion. Thereafter, eluted peptides were lyophilized and re-suspended in 5% acetonitrile, 0.1% TFA. Each sample was loaded onto a 25□cm × 75□μm ID, 3 μm, 100Å Magic C18AQ analytical column (Acclaim, Thermo Scientific). Peptides were eluted into Orbitrap QExactive HF-X mass spectrometer (ThermoFisher Scientific) in 120 min using a 5%–35% linear gradient of acetonitrile in 0.1% formic acid and water with 0.1% formic acid. The full scan was acquired with a resolution of 70,000 resolution (at *m/z* 200). Afterwards, ten most abundant data-dependent MS/MS spectra were collected with a resolution of 17,500 (at *m/z* 200), AGC target of 1 × 10^6^, a maximum ion time of 120 ms, and normalized collision energy of 27.

### Isolation of mouse germ cells by fluorescence-activated cell sorting

Male germ cell sorting was performed as previously described ^22^. Briefly, testes were decapsulated and incubated in 1× Gey’s Balanced Salt Solution (GBSS, Sigma, G9779) supplemented with 0.4 mg/mL collagenase type IV (Worthington, LS004188) at 33°C for 15 min. Following two washes in 1× GBSS, seminiferous tubules were incubated in 1× GBSS containing 0.5 mg/mL trypsin and 1 μg/mL DNase I at 33°C for 15 min. The tubules were then placed on ice at 4°C and mechanically dissociated by gentle Pasteur pipetting for 3 min. Trypsin activity was quenched by adding fetal bovine serum (FBS) to a final concentration of 7.5% (v/v), after which the cell suspension was passed through a pre-wetted 70 μm cell strainer. Cells were pelleted by centrifugation at 300 × g for 10 min at 4°C and resuspended in 1× GBSS containing 5% (v/v) FBS, 1 μg/mL DNase I, and 5 μg/mL Hoechst 33342 (Thermo Fisher, 62249). The suspension was incubated at 33°C for 45 min using a rotator at 150 rpm. Propidium iodide (PI) was then added at a final concentration of 0.2 μg/mL (Thermo Fisher, P3566), and the cells were passed through a pre-wetted 40 μm cell strainer. Four-way cell sorting was performed using a BD FACSMelody Cell Sorter (BD Biosciences) to isolate spermatogonia (SpG), leptotene and zygotene (L/Z), pachytene and diplotene (P/D), and round spermatids (RS) population according to our previously established gating strategies ^22^. The 488-nm laser was used to excite PI whose emission was detected through a 700/54 nm bandpass filter. Hoechst 33342 was excited with a 405-nm laser, and its emission was recorded using 660/10-nm (red) and 448/45-nm (blue) bandpass filters.

### Chromosome spread

Chromosome spreads were prepared as previously described^22^. Briefly, cells were incubated in 25 mM sucrose solution at room temperature for 20 min and fixed in 1% (w/v) paraformaldehyde solution containing 0.15% (v/v) Triton X-100 at room temperature for 2 h. Fixed cells were then added dropwise onto SuperFrost Plus microscope slides (VWR, 631-108), which were pre-dipped in the same solution, and incubated in a humidifying chamber for at least 2 h before air drying. The dried slides were washed sequentially at room temperature for 10 min with the following solutions: (i) 1× PBS containing 0.4% (v/v) Photo-Flo 200 (Kodak, 1026269), (ii) 1× PBS containing 0.1% (v/v) Triton X-100, and (iii) ADB/PBS solution, which consists of 10% ADB [0.03% (w/v) BSA, 0.01% (v/v) Triton X-100, 10% (v/v) donkey serum (Sigma, D9663), 2× PBS] mixed with 90% 1× PBS. Slides were then incubated overnight at 4°C with primary antibodies diluted in ADB solution as follows: anti-γH2AX (Novus Biologicals, NB100-384, 1:1,000), and anti-SCP-3 (D-1) (Santa Cruz, sc-74569, 1:400). The next day, slides were washed as described above and incubated at 37°C for 1 h with Alexa Fluor anti-mouse 594 (Thermo Fisher Scientific, A21203, 1:2,000) and Alexa Fluor anti-rabbit 488 (Thermo Fisher Scientific, A21206, 1:2,000) secondary antibodies. After three 10-min washes in 1× PBS containing 0.4% (v/v) Photo-Flo 200 and one 10-min wash in 0.4% (v/v) Photo-Flo 200 alone, slides were dried in the dark and mounted with ProLong Gold Antifade Mountant with DAPI (Thermo Fisher Scientific, P36931). Images were captured using a Zeiss Axio Observer 7 inverted widefield microscope.

### Testis histology by hematoxylin and eosin staining

Testis tissues were fixed in Bouińs solution (VWR, 7000.1000). After 24 h at room temperature, fixed tissues were transferred to 70% ethanol, embedded in paraffin blocks, and sectioned at a thickness of 5 µm. The sections were stained using hematoxylin and eosin (H&E) staining kit according to the manufacturer’s protocol (Abcam, ab245880). Images of H&E staining were captured using a Zeiss Axio Observer 7 inverted widefield microscope.

### Immunofluorescence staining

Tissue sections were deparaffinized by sequential incubation in xylene (7 minutes × 3), followed by ethanol washes (100%, 95%, and 70% (v/v) ethanol). Antigen retrieval was performed by boiling the slides in 1× citrate buffer (pH 6.0) at 95–100°C for 20 min, followed by cooling at room temperature for 10 min and a single wash with PBS. Slides were placed in a humidified chamber, and the tissue area was outlined with a PAP pen (Thermo Fischer Scientific, 008899). After three washes with PBS (5 min each), sections were blocked with 5% (w/v) bovine serum albumin (Sigma, A9418) at room temperature for 1 h. Samples were then incubated overnight at 4°C with primary antibodies diluted in 1 × PBS: anti-SMARCA5 (Thermo Fisher Scientific, PA5-52601, 1:200), anti-DDX4/MVH (Abcam, ab13840, 1:500), anti-γH2AX (Novus Biologicals, NB100-384, 1:2,000), FITC-conjugated peanut agglutinin (Sigma, L7381, 1:1,000), anti-HSPA2 (Sigma, HPA000798, 1:500), anti-SALL4 (Abcam, ab57577, 1:1,000), anti-c-KIT (Abcam, ab317843, 1:250), anti-STRA8 (Proteintech, CL488-68071, 1:500), anti-H3S10P (Abcam, ab14955, 1:500), anti-GATA4 (eBiosciences, 14998080, 1:100), and anti-SOX9 (Sigma, ABE571, 1:700). Following primary antibody incubation, slides were washed three times with PBS containing 0.02% Tween-20 for 10 min each. Secondary antibodies were then applied at room temperature for 1 h: anti-rabbit Alexa Fluor 488 (Thermo Fisher Scientific, A21206, 1:700), anti-rabbit Alexa Fluor 594 (Thermo Fisher Scientific, A211012, 1:700), anti-mouse Alexa Fluor 594 (Thermo Fisher Scientific, A21203, 1:700), anti-mouse Alexa Fluor 488 (Thermo Fisher Scientific, A21202, 1:700), anti-mouse Alexa Fluor 647 (Thermo Fisher Scientific, A21235, 1:200), and anti-rat Alexa Fluor 594 (Thermo Fisher Scientific, A21209, 1:700). Nuclei were counterstained with DAPI (Sigma, D1306, 1:1,000) for 10 min. Finally, slides were washed twice with PBS for 10 min each and mounted with coverslips using DABCO mounting media (Sigma, D27802). Immunoflourescence images were acquired by Zeiss Axio Observer 7 inverted widefield microscope.

### TUNEL staining

The TdT-mediated dUTP Nick-End Labeling (TUNEL) assay was performed using the DeadEnd Fluorometric TUNEL System (Promega, G3250) according to the manufacturer’s instructions. Tissue sections at 5 μm thickness were deparaffinized by immersing twice in xylene for 5 min each, followed by incubation in 100% ethanol at room temperature for 5 min. The sections were then rehydrated through sequential incubations in descending ethanol concentrations for 3 min each: 100%, 95%, 85%, 70%, and 50% (v/v). Following rehydration, the slides were washed once in 0.85% (w/v) NaCl and once in 1× PBS at room temperature for 5 min each. The tissue sections were then fixed in 1× PBS containing 4% (w/v) paraformaldehyde at room temperature for 15 min. Thereafter, slides were washed twice in 1× PBS for 5 min each, and treated with 20 µg/ml proteinase K at room temperature for 9 min. After proteinase K treatment, sections were washed once in 1× PBS at room temperature for 5 min and fixed again with 1× PBS containing 4% (w/v) paraformaldehyde at room temperature for 5 min. Excess liquid was removed from the slides, and the tissue sections were covered with 100 μl of Equilibration Buffer at room temperature for 5 min. The tissue sections were then incubated with 50 μl rTdT incubation buffer (45 μl Equilibration Buffer, 5 μl Nucleotide Mix containing fluorescein-12-dUTP, and 1 μl rTdT Enzyme) in a humidifying chamber at 37°C for 1 h. To terminate the reaction, the slides were immersed in 2× saline-sodium citrate (SSC) buffer for 15 min, followed by three sequential washes in 1× PBS for 5 min each. Thereafter, the tissue sections were stained in 1× PBS containing 1 μg/ml propidium iodide at room temperature for 15 min in the dark. Finally, the slides were washed three times in deionized water for 5 min each, mounted with one drop of DABCO (Sigma, D27802), and covered with a glass coverslip. Images were captured using a Zeiss Axio Observer 7 inverted widefield microscope.

### Western blotting

Finely minced tissue samples and cell lines were homogenized in a Dounce homogenizer using 20–30 strokes of pestle B in RIPA lysis buffer (Thermo Fisher Scientific, 89900, 25 mM Tris-HCl, pH 7.6, 150 mM NaCl, 1% (v/v) NP-40, 1% (w/v) sodium deoxycholate, and 0.1% (w/v) SDS) supplemented with 1% (v/v) protease inhibitor (Thermo Fisher Scientific, 1861279). Lysates were incubated on ice for 30 min and were subsequently centrifuged at 20,000g at 4°C for 30 min. The supernatant was transferred to a fresh tube, and protein concentrations were measured using the BCA Protein Assay Kit (Thermo Fisher Scientific, 23227). A total of 75 μg protein was mixed with one-fourth volume of loading dye (Invitrogen, NP0007) containing 0.2M dithiothreitol (DTT) and heated at 95°C for 5 min to denature proteins. The protein samples were resolved on 4–20% Tris-Glycine gradient SDS gel (Thermo Fisher Scientific, XP04200) and transferred to a 0.45 µm nitrocellulose membrane (Thermo Fisher Scientific, 88018). Membranes were blocked using Blocking Buffer (Thermo Fisher Scientific, 37565) at room temperature for 1.5 h and incubated with primary antibodies at 4°C overnight: rabbit anti-SMARCA5 (Atlas, HPA00875, 1:750), rabbit anti-SMARCA5 (Abcam, ab3749, 1:500), mouse anti-NFYA (Santa Cruz Biotechnology, sc-17753, 1:500), and mouse anti-ACTIN (Sigma, A1978, 1:5,000). Next day, the membranes were washed three times with 1× PBS-T (0.1% (v/v) Tween-20 in 1× PBS) at room temperature 30 min each. Membranes were then incubated with secondary antibodies at room temperature for 1h: anti-rabbit IRDye 800CW (LI-COR Biosciences, 926–32213, 1:15,000), and anti-mouse IRDye 680RD (LI-COR Biosciences, 926–68072, 1:15,000), followed by three washes with 1× PBS-T. Protein signal was detected using the Odyssey Infrared Imaging System (LI-COR).

### Co-immunoprecipitation of NFYA and SMARCA5 in HEK293T cells

HEK293T cells were seeded at a density of 8 × 10^5^ cells in 6-cm dishes one day prior to transfection. Cells were transfected using the TransIT-X2® Dynamic Delivery System (Mirus Bio, MIR 6000) and Opti-MEM I Reduced Serum Medium (Thermo Fisher Scientific, 31985062) according to the manufacturer’s instructions. A total of 5.5 μg plasmid DNA was used under two conditions: (1) pCMV3-N-FLAG-NFYA (Sino Biological, HG18080-NF) co-transfected with pCMV3-HA-SMARCA5 (Sino Biological HG18406-NY); and (2) pCMV3-HA-SMARCA5 alone. 48 h after transfection, cells were fixed using 0.5mM dithiobis(maleimido)ethane (DTME) (Thermo Fisher Scientific, 22335) and incubated for 15 min on a shaker. Cells were subsequently harvested in RIPA buffer (Thermo Fisher Scientific, 89900, 25 mM Tris-HCl, pH 7.6, 150 mM NaCl, 1% (v/v) NP-40, 1% (w/v) sodium deoxycholate, and 0.1% (w/v) SDS) supplemented with 1% protease inhibitor (Thermo Fisher Scientific, 1861279). 10% of the total lysate was flash-frozen and stored at −80□°C as input, while the remaining lysate was incubated overnight at 4□°C with rotation in the presence of 40 μl anti-FLAG M2 paramagnetic beads (Sigma, M8823). The following day, beads were washed five times for 5□min each with RIPA buffer containing 1% protease inhibitor at 4□°C. Bound proteins were eluted using 50 μl 3×FLAG peptide (Sigma, F4799) for 2.5□h at room temperature with rotation. Eluted proteins were subsequently analyzed by western blotting using rabbit anti-HA (H6908, Sigma, 1:3,000) and mouse anti-FLAG (Sigma, f1804, 1:3,000) primary antibodies.

### Luciferase assay

One day before transfection, GAL4 reporter HEK293T cells (BPS Bioscience, 60656) were seeded at a density of 3 × 10□cells per well in 96-well plates. Cells were transfected with GAL4-empty (VectorBuilder, VB230515-1632PWZ), HA-SMARCA5 (Sino Biological, HG18406-NY), FLAG-NFYA (Sino Biological, HG18080-NF), GAL4-SMARCA5 (VectorBuilder, VB230515-1203MDB), GAL4-NFYA (VectorBuilder, VB230514-1234KQS), or GAL4-NFYA together with HA-SMARCA5 using TransIT-X2® Dynamic Delivery System (Mirus Bio, MIR 6000) according to the manufacturer’s instructions. pRL-TK Renilla luciferase control plasmid (Promega, E2241) was co-transfected for normalization. Luciferase activity was measured 48 h after transfection using the Dual-Glo® Luciferase Assay System (Promega, E2920) according to the manufacturer’s instructions. The experiment was repeated independently 10 times, each with technical triplicates.

### CUT&Tag Sequencing

<u>C</u>leavage <u>U</u>nder <u>T</u>argets and <u>Tag</u>mentation (CUT&Tag) protocol was adopted from ^85^ (DOI: dx.doi.org/10.17504/protocols.io.x54v9mkmzg3e/v5). Cells were washed once in PBS and incubated in NE1 buffer (0.02 M [w/v] HEPES-KOH pH 7.9, 0.01 M [w/v] KCl, 0.5 mM [w/v] Spermidine, 0.1% [v/v] TritonX-100, 10% [v/v] Glycerol, 1 × protease inhibitor cocktail [PIC]) on ice for 10 min. After centrifuging at 1300 × g at 4°C for 4 min, nuclei were resuspended in PBS. Isolated nuclei were crosslinked with 0.1% formaldehyde at room temperature for 2 min. Crosslinking was stopped by the addition of glycine to a final concentration twice that of formaldehyde. Cells were centrifuged again as previously and resuspended in Wash buffer (0.02 M [w/v] HEPES pH 7.5, 0.15 M [w/v] NaCl, 0.5 mM [w/v] Spermidine, 1 × PIC).

80,000 fixed nuclei or unfixed cells were incubated with 5 μl Concanavalin A-coated magnetic beads (Cell Signaling, 82307S) at room temperature for 10 min. After removing the supernatant, the beads were resuspended in 40 μl antibody buffer (20 mM (w/v) HEPES (pH 7.5), 150 mM (w/v) NaCl, 0.5 mM (w/v) spermidine, 1% (v/v) Triton X-100, 1× PIC, 0.1% (w/v) BSA, 2 mM (w/v) EDTA) for separate samples containing 0.5 µg anti-SMARCA5 (EpiCypher, 13-2007) or 0.5 µg anti-BPTF antibody (Sigma, ABE1966) at a 1:50 dilution. The beads were then gently resuspended by mild vortexing. Following the incubation with the primary antibody at room temperature for 1 h, the supernatant was discarded and the cell–bead mixture was resuspended in 40 μl Triton-wash buffer (20 mM (w/v) HEPES (pH 7.5), 150 mM (w/v) NaCl, 0.5 mM (w/v) spermidine, 1% (v/v) Triton X-100, and 1× PIC), supplemented with guinea pig anti-rabbit secondary antibody (Antibodies Online, ABIN101961) at a 1:100 dilution. Samples were then incubated at room temperature for 30 min. The cells were washed once with 200 μl Triton-wash buffer before being resuspended in 40 μl Triton-300 wash (20 mM (w/v) HEPES pH 7.5, 300 mM (w/v) NaCl, 0.5 mM (w/v) Spermidine, 1% (v/v) Triton-X100, 1 x PIC) buffer containing preloaded pAG-Tn5 fusion protein at a 1:25 dilution and incubated at room temperature for 1 h. Following pAG-Tn5 binding, the cells were washed once with 200 μl Triton-300 wash buffer and subsequently resuspended in 50 μl CUTAC-DMF tagmentation solution containing 10 mM (w/v) TAPS (pH 8.5), 5 mM (w/v) MgCl_₂_, and 20% (v/v) DMF. Thereafter, the samples were incubated in a thermocycler at 55°C for 1 h. After tagmentation, the cells were washed once with 50 μl TAPS wash buffer (10 mM (w/v) TAPS (pH 8.5) and 0.2 mM (w/v) EDTA) followed by resuspension in 5 μl 1% SDS-ProtK release solution (1% (w/v) SDS and 10 mM (w/v) TAPS (pH 8.5), Thermolabile Proteinase K (NEB, P8111S) at a 1:11 dilution). The samples were thoroughly mixed and incubated in a thermal cycler at 37°C for 1 h, followed by an additional incubation at 58°C for 1 h. Following incubation, 15 μl 6% Triton X-100 together with 2 μl of 10 μM i7 indexed primer and 2 μl 10 μM i5 indexed primer were added to the samples. The supernatant was subsequently transferred to a fresh PCR tube, mixed with 25 μl NEBNext 2× PCR Master Mix (NEB, M0541L), and amplified in a thermal cycler using the following conditions: 58°C for 5 min, 72°C for 5 min, and 98°C for 5 min, followed by 11 cycles of 98°C for 10 s and 60°C for 10 s, with a final extension at 72°C for 1 min. Following PCR amplification, libraries were purified using 65 μl AMPure XP beads (Beckman Coulter, A63880) and eluted in 22 μl Tris-HCl (pH 8.5). CUT&Tag libraries were sequenced on a NextSeq550 system (Illumina) using 79 + 79 nt paired-end reads.

### Simultaneous single-cell ATAC-seq and mRNA-seq profiling

We used the BD Rhapsody Single-Cell Analysis System to simultaneously examine the chromatin accessibility and mRNA expression profile in the testicular cells of 8-week-old *Smarca5-CKO* mouse.

#### Isolation of nuclei for single-cell multiomics

Testis was dissected and gently dispersed in 6 ml of 1× Gey’s Balanced Salt Solution (GBSS, Sigma, G9779) supplemented with 0.4 mg/ml collagenase type IV (Worthington, LS004188), followed by incubation at 33°C with shaking at 600 rpm for 15 min. The seminiferous tubules were subsequently washed twice with 6 ml GBSS and further incubated in 1× GBSS containing 0.5 mg/ml trypsin (Gibco, 27250-018) and 1 µg/ml DNase I (Roche, 10104159001) at 33°C and 600 rpm for 15 min. The tubules were subsequently homogenized using a Pasteur pipette on ice at 4°C for 3 min. To inactivate trypsin, 400 μl FBS was added, and the cell suspension was filtered through a pre-wetted 70 μm cell strainer (Corning, CLS431751). Cells were then pelleted by centrifugation in a swing-bucket centrifuge at 500 × g for 10 min at 4°C and resuspended in 3.5 ml 1× GBSS supplemented with 175 μl FBS and 6 μl DNase I. Following resuspension, the cell suspension was passed through a pre-wetted 40 μm cell strainer (Corning, 43750). A total of 300,000 cells were collected for nuclei isolation and pelleted by centrifugation at 400 × g at 4°C for 10 min. The cells were then gently resuspended in 195 μl swelling buffer containing 10 mM (w/v) Tris-HCl (pH 7.5), 2 mM (w/v) MgCl_₂_, and 3 mM (w/v) CaCl_₂_. Dropwise, an additional 1.8 ml of swelling buffer was applied to the resuspended cells. After 5 min of ice incubation, the cells were pelleted at 400 × g at 4°C for 10 min. After careful removal of the supernatant, the cell pellet was resuspended in 100.5 μl pre-chilled lysis buffer 1 (9 mM (w/v) Tris-HCl pH 7.5, 1.8 mM (w/v) MgCl2, 2.7 mM (w/v) CaCl2, 10% (v/v) glycerol, 1 x protease inhibitor cocktail (PIC), and 0.4 U/μl RNasin (Promega, N2615). Next, the suspension was dropwise mixed with 100.5 μl of lysis buffer 2 (9 mM (w/v) Tris-HCl (pH 7.5), 1.8 mM (w/v) MgCl_₂_, 2.7 mM (w/v) CaCl_₂_, 10% (v/v) glycerol, 10% (v/v) Igepal CA-360, 1× protease inhibitor cocktail (PIC), and 0.4 U/μl RNasin (Promega, N2615). The mixture was then kept on ice for 5 min to allow cell lysis. The isolated nuclei were collected by centrifugation at 600 × g at 4°C for 5 min. Following removal of the supernatant, the nuclei pellet was gently resuspended in 200.25 μl lysis buffer 3 (9 mM (w/v) Tris-HCl (pH 7.5), 1.8 mM (w/v) MgCl_₂_, 2.7 mM (w/v) CaCl_₂_, 10% (v/v) glycerol, 0.01% (v/v) digitonin, 1× protease inhibitor cocktail (PIC), and 0.4 U/μl RNasin [Promega; N2615]). An additional 1.8 ml lysis buffer 3 was then gradually added to the nuclei suspension with gentle mixing. The nuclei were subsequently pelleted by centrifugation at 500 × g at 4°C for 5 min. After discarding the supernatant, the nuclei pellet was resuspended in 200.25 μl freezing buffer (50 mM (w/v) Tris-HCl (pH 8), 5 mM (w/v) MgCl_₂_, 40% (v/v) glycerol, 1× protease inhibitor cocktail (PIC), and 0.4 U/μl RNasin (Promega, N2615). The nuclei were pelleted by centrifugation at 900 × g at 4°C for 6 min, after which the supernatant was carefully removed. The nuclei pellet was then resuspended in 50 μl modified nuclei buffer (BD Biosciences, 51-9023091) supplemented with 2.5% (v/v) RNase inhibitor (BD Biosciences, 51-9024039) and 1 mM DTT. Subsequently, 50,000 nuclei were used for preparation of ATAC-seq and mRNA-seq libraries.

### Simultaneous ATAC-seq and mRNA whole transcriptome (WTA) library preparation

For DNA tagmentation, 50,000 nuclei suspended in 5 μl modified nuclei buffer (BD Biosciences, 667052) were combined with 25 μl Tagmentation buffer with 20% DMF, 11.75 μl water, 2 μl 10× PBS, 1.25 μl RNase inhibitor (BD Biosciences, 51-9024039), 0.5 μl 1% (v/v) digitonin, 0.5 μl 10% (v/v) Tween-20, and 4 μl Tagmentase. The reaction mixture was incubated at 37°C for 30 min. Following incubation, 300 μl modified sample buffer was added, and the nuclei suspension was subsequently loaded onto a BD Rhapsody 8-Lane Cartridge (BD Biosciences, 666262). The cartridge was first incubated at room temperature for 8 min to facilitate settling of nuclei into the cartridge microwells. The nuclei were subsequently washed twice with cold sample buffer and incubated with splint beads (BD Biosciences, 667052) at room temperature for 3 min. Afterwards, the cartridge was briefly shaken for 10 s at 1,000 rpm at room temperature using a ThermoMixer (Eppendorf, EP5382000015), followed by two additional washes with cold sample buffer.

Subsequently, 280 μl lysis buffer containing 5 mM DTT and 25 μl proteinase K was added to the cartridge, followed by incubation at 37°C for 10 min. Following lysis, the tagmented DNA and mRNA were fixed onto beads using splint-oligo-bonded TSO and poly(T) oligo, respectively. The beads carrying the captured tagmented DNA and mRNA were subsequently transferred into a 1.7 ml tube and washed for two times with cold Bead wash buffer. Tagmented DNA fragments were then ligated to the BD Rhapsody bead oligos in a ligation reaction containing 20 μl 10× ligation buffer, 10 μl DNA ligase (BD Biosciences, 41928), 5 μl RNase inhibitor (BD Biosciences, 51-9024039), and 170 μl water. The reaction was incubated at 25°C for 30 min with shaking at 1200 rpm. Following ligation, the ligation mixture was removed and reverse transcription (RT) was carried out in 200 μl RT reaction mix containing 40 μl RT buffer (BD Biosciences, 633773), 20 μl 10 mM dNTPs, 10 μl 0.1 M DTT, 12 μl RT/PCR enhancer, 10 μl RNase inhibitor (BD Biosciences, 633773), 10 μl reverse transcriptase (BD Biosciences, 633773), and 98 μl water. The reaction was incubated at 42°C for 30 min with shaking at 1200 rpm. After completion of the RT reaction, the RT mixture was exchanged with splint-oligo removal buffer, and the samples were incubated at 60°C for 5 min with shaking at 1200 rpm, then 5 min on ice. To eliminate excess oligos remaining on the beads, the samples were subsequently treated with 200 μl Exonuclease I reaction mix containing 20 μl 10× Exonuclease I buffer, 10 μl Exonuclease I (BD Biosciences, 633773), and 170 μl water, followed by incubation at 37°C for 30 min at 1200 rpm. Exonuclease I activity was subsequently terminated by addition of 4 μl 0.5 M EDTA to the reaction mixture.

For ATAC-seq library preparation, DNA was eluted from the beads twice using 40 μl elution buffer per round. Following DNA elution, the beads were preserved in bead resuspension buffer at 4°C for subsequent whole-transcriptome amplification (WTA) library preparation. The 80 μl ATAC-seq libraries were then amplified and indexed in a 42 μl PCR indexing reaction containing 30 μl PCR master mix (BD Biosciences, 41928), 6 μl ATAC-seq forward primer and 6 μl ATAC-seq reverse primer. PCR amplification was performed under the following cycling conditions: 98°C for 45 s, followed by 12 cycles of 98°C for 10 s, 66°C for 30 s, and 72°C for 30 s, with a final extension step at 72°C for 1 min. For WTA library preparation, the beads stored at 4°C were initially heated at 95°C for 5 min. The beads were then separated using a magnet, the supernatant was removed, and the beads were resuspended in 87 μl Random Primer mix containing 10 μl WTA extension buffer (BD Biosciences, 633801), 10 μl WTA extension primer, and 67 μl water. The samples were incubated at 95°C for 5 min, followed by incubation at 37°C and 25°C for 5 min each with shaking at 1200 rpm. Subsequently, 13 μl extension enzyme mix consisting of 8 μl 10 mM dNTPs, 12 μl ET/PCR enhancer, and 6 μl WTA extension enzyme (BD Biosciences, 633801) was added to the reaction. The mixture was then incubated in a ThermoMixer (Eppendorf, EP5382000015) at 1200 rpm using the following program: 25°C for 10 min, 37°C for 15 min, 45°C for 10 min, and 55°C for 10 min. To recover the Random Primer Extension (RPE) products from the beads, the samples were incubated in 205 μl elution buffer at 95°C for 5 min. The WTA random priming, extension, and purification steps were subsequently repeated once more. In total, 400 μl RPE product was purified using 1.8× AMPure XP beads (Beckman Coulter, A63881) and finally eluted in 40 μl elution buffer. Following elution, 60 μl PCR master mix supplemented with 10 μl Universal oligo and 10 μl WTA amplification primer was directly combined with the 40 μl eluted RPE product for the initial amplification of WTA libraries. PCR amplification was carried out under the following conditions: 95°C for 3 min, followed by 11 cycles of 95°C for 30 s, 60°C for 1 min, and 72°C for 1 min, with a final extension at 72°C for 2 min. The amplified products were subsequently purified using 0.8× AMPure XP beads (Beckman Coulter, A63881). Prior to the final amplification step, WTA libraries were evaluated using Agilent 2100 Bioanalyzer to determine the molar concentration of DNA fragments within the 150–600 bp size range. Subsequently, 10 μl of 2 nM WTA library was subjected to indexing in a 40 μl PCR indexing reaction containing 25 μl PCR master mix (BD Biosciences, 633801), 5 μl BD Rhapsody library forward primer, 5 μl BD Rhapsody reverse indexed primer, and 5 μl water. PCR amplification was performed under the following cycling conditions: 95°C for 3 min, followed by 8 cycles of 95°C for 30 s, 60°C for 30 s, and 72°C for 30 s, with a final extension at 72°C for 1 min. ATAC-seq and WTA libraries were sequenced on the NovaSeq X Plus platform (Illumina) using the following run configurations: ATAC-seq libraries were sequenced with 50 cycles for Read 1, 50 cycles for Read 2, 8 cycles for Index 1, and 60 cycles for Index 2, whereas WTA libraries were sequenced using 51 cycles for Read 1, 71 cycles for Read 2, and 8 cycles for Index 1. Libraries were sequenced according to the manufacturer’s recommended depth, corresponding to 50,000 read pairs per cell for ATAC-seq libraries and 100,000 read pairs per cell for WTA libraries.

### CUT&RUN and CUT&Tag analyses

Along with our SMARCA5 and BPTF CUT&Tag datasets from FACS-purified SpG, we re-analyzed publicly available H2AK119ub CUT&Tag^51^ (GSE221942) and NFYA CUT&RUN^22^ (GSE295991) datasets FACS-purified SpG. Adapter sequences and low-quality bases were trimmed from the raw paired-end reads using Fastp (v1.3.4^78^). Reads were aligned to the mouse reference genome (mm10) using Bowtie2 (v2.5.4^69^) with the parameters —local —very-sensitive —no-unal —no-mixed —no-discordant-|10-X700. Aligned reads were converted from SAM to BAM format, sorted and indexed using SAMtools (v1.21 ^74^). Thereafter, duplicate reads were removed using MarkDuplicates program from picard-tools (v.3.4.0). Only uniquely mapped reads were used for downstream analysis. Genome-wide coverage BedGraph files were generated using BEDTools (v2.31.1^76^) genomecov (-bg) and restricted to intervals with non-zero coverage. Significant peaks in CUT&Tag experiments were called using SEACR v1.3^71^ with the following settings −0.01-non-stringent, whereas peaks of CUT&RUN experiments were called using MACS3 (v3.0.2^70^; FDR < 0.1). IgG antibody was used as background to call significant peaks of CUT&RUN and CUT&Tag experiments. Genes with a factor peak within ±2□kb of their transcription start sites in at least two biological replicates of CUT&RUN or CUT&Tag were considered as factor-bound genes.

### ChIP-seq analysis

We re-analyzed publicly available SALL4 and ZBTB16 ChIP-seq from testis of seven days postpartum^23^ (GSE202819) and H3K27Ac ChIP-seq from FACS-purified differentiating SpG^50^ (GSE132446). Data analysis was performed as described for CUT&RUN and CUT&Tag.

## Simultaneous single-cell ATAC-seq and mRNA-seq analysis

### scRNA-seq analysis

#### Cell barcode and UMI assignment

Valid cell barcodes were obtained from BD Biosciences. Reads lacking a valid cell barcode were discarded, and the cell barcode and unique molecular identifier (UMI) of the remaining reads were assigned to their corresponding cDNA reads using BD Rhapsody Sequence Analysis Pipeline (v3.0^72^). These barcode- and UMI-tagged reads were used to generate gene-expression count matrices for each sample.

#### Read alignment and count matrix generation

Reads were aligned to the mouse reference genome (mm10) using STAR (v2.2.7b^73^) and assigned to annotated genes (GENCODE M25). PCR duplicates sharing the same cell barcode, UMI and alignment coordinates were removed, and the UMI-collapsed counts were used to build a sparse cell-by-gene count matrix for each sample. The count matrices were then imported into Seurat for downstream analysis.

#### Quality control and doublet removal

Data were analyzed with the Seurat R package (v5.2.1^82^). Cells with fewer than 500 detected genes, or with ≥5% of UMIs derived from mitochondrial genes, were removed. Doublets and multiplets were identified in each sample separately using DoubletFinder (v2.0.4^83^) with the top 10 principal components (PCs) and default parameters, and cells classified as doublets were excluded before the samples were combined.

#### Normalization, dimensionality reduction and clustering

Filtered singlets from the *Cre/+*, *Nfya-CKO* and *Smarca5-CKO* mice were merged into a single Seurat object, and the RNA assay layers were joined. Gene UMI counts were log-normalized with a scale factor of 10,000, and the top 5,000 variable genes were identified, scaled and used for principal component analysis (PCA). The top 30 PCs were used to build the nearest-neighbor graph, which was clustered at a resolution of 0.8, and to compute a uniform manifold approximation and projection (UMAP). After quality control and doublet removal, 78,860 cells (67,549 cells from *Cre/+*, 5,061 cells from *Nfya-CKO* and 6,250 cells from *Smarca5-CKO*) were grouped into 33 clusters.

#### Cell type annotation

Based on the expression of established marker genes, the 33 clusters were assigned to 13 major cell populations, including successive germ-cell populations spanning the spermatogenic trajectory from undifferentiated spermatogonia to elongating spermatids, together with the major somatic cell populations. Clusters 7 and 22 could not be confidently assigned and were labeled as unknown.

#### Differential gene expression analysis

Differentially expressed genes between selected cell populations were identified from RNA expression data using Seurat’s FindMarkers function with the Wilcoxon rank-sum test.

#### scATAC-seq analysis

Raw scATAC-seq reads were processed using the BD Rhapsody Sequence Analysis Pipeline (v3.0^72^). Following initial quality control and removal of low-quality reads, reads were aligned to the mouse reference genome (mm10) using bwa-mem2 (v2.2.1^81^). The processed scATAC-seq data were then imported into ArchR (v1.0.3^62^) for quality control, dimensionality reduction, clustering, and downstream chromatin accessibility analysis. Cells with fewer than 1,000 fragments or a transcription start site (TSS) enrichment score below 4 were removed, and putative doublets were identified and excluded using ArchR’s doublet detection. After quality control and doublet removal, 64,361 cells remained for downstream analysis (55,952 cells from *Cre/+*, 4,258 cells from *Nfya-CKO,* and 4,151 *Smarca5-CKO* mice), with a mean TSS enrichment score of 10.15 (Supplemental Table S6). Dimensionality reduction and clustering were performed on a genome-wide matrix of 500-bp binarized tiles (TileMatrix). We applied iterative latent semantic indexing (LSI) to this matrix and visualized the result by UMAP. Cell identities were assigned by combining annotations transferred from the corresponding scRNA-seq data with chromatin accessibility at established cell-type-specific marker genes. The resulting UMAP resolved the major somatic and germ-cell populations and reflected the developmental progression of spermatogenesis.

To visualize and quantify chromatin accessibility, cells within each annotated population were pooled into pseudobulk profiles. bigWig tracks were generated in ArchR at 100-bp resolution using nFrags normalization, in which the pooled signal was scaled by 10^4 relative to the total fragment count of each cell group. TSS-centered accessibility was quantified from these nFrags-normalized pseudobulk tracks. Signal was summarized in 100-bp bins within ±2 kb of annotated gene TSSs.

## Quantification and statistical analysis

Statistical analysis and graph plotting was performed in R 4.6.1 (https://www.rstudio.com) and Prism v 11.0.1 (GraphPad Software, LLC). Box or violin plots were used to represent data distribution. The boxes represent the interquartile ranges (IQRs). Data points with values > third quartile + 1.5 × IQR or lower than the first quartile − 1.5 × IQR were defined as outliers. IQR is defined as the difference between the minimum of the first quartile and the maximum of the third quartile. Spearman’s correlation (*p*) was used to measure relationship between variables. ANOVA test was applied to measure the significant differences between conditions in Figure 1A. Two-sided paired Student *t* test was used to calculate the significance in Figures 1D, 2B, 2D-F and Document S1: Figures S3D, S3E, S4B, S5A-D. Two-sided wilcoxon matched-pairs signed-ranked sum test used in Figures 1E, 4E, 4G, 6C, 6D. Mann–Whitney–Wilcoxon *U* test was used to calculate significance in Figures 4D, 7A-B.

