## Supplemental Data 1 for "Active maintenance of accessible chromatin at poised meiotic promoters by the NFYA-NURF axis is required for meiotic entry in mice"

**Figure S1. NFYA interacts with SMARCA5 in mouse testes and heterologous cells. Related to Figure 1.**

**(A)** Western blotting shows the abundance of NFYA in the eluate fractions from NFYA and IgG immunoprecipitations in the testis and liver extracts of adult mice. IgG immunoprecipitations served as negative controls. Dashed rectangle indicates the region presented Figure 1B. In input, FT flow-through, E eluate.

**(B)** Western blotting shows the abundance of FLAG-tagged NFYA and HA-tagged SMARCA5 proteins upon co-immunoprecipitation of FLAG-NFYA and HA-SMARCA5 in HEK293. Cells expressing HA-SMARCA5 without FLAG-NFYA served as negative control. Dashed rectangle indicates the region presented in Figure 1C. In input, FT flow-through, E eluate.

**(C)** Western blotting shows the abundances of NFYA and SMARCA5 upon co-immunoprecipitation of endogenous SMARCA5 with NFYA from adult mouse testis extracts. IgG immunoprecipitation served as a negative control. Dashed rectangle indicates the regions presented in Figure 1E. In, input; FT, flow-through; W1, first wash; E, eluate.

**Figure S2. SMARCA5 expression is enriched in pre-meiotic germ cells. Related to Figure 2.**

**(A)** Violin plots show the mRNA abundances of *Smarca5* (left) and *Bptf* (right) across developing male germ cells using our previously published single-cell RNA-seq dataset^22^. SpG, spermatogonia; pL, preleptotene; eL, early leptotene; L, leptotene; Z, zygotene; eP, early pachytene; mP, middle pachytene; lP, late pachytene; D, diplotene; RS, round spermatids; ES, elongating spermatids.

**(B)** Immunofluorescence staining for SMARCA5 in the testes sections from wild-type mice at 9, 14, 17, and 21 days post-partum (dpp). SpG, spermatogonia; A, type A spermatogonia; In, intermediate spermatogonia; B, type B spermatogonia; L, leptotene; L/Z, leptotene/zygotene; eP, early pachytene; P, pachytene; RS, round spermatids. Scale bars: 50 µm.

**Figure S3. Generation and validation of germ cell-specific Smarca5 knockout mice. Related to Figure 2.**

**(A)** Schematic representation of the strategy used to generate germ cell-specific *Smarca5* knockout mice using the Stra8-Cre-loxP system. Exons 5 and 6 of *Smarca5* were flanked by loxP sites and excised in pre-meiotic germ cells expressing the *Stra8-P2A-ZsGreen1-T2A-Cre* transgene.

**(B)** Images for the agarose gel electrophoresis showing PCR genotyping of *Smarca5* floxed alleles and the *Stra8-Cre* transgene. PCR-1 and PCR-2 were used to validate the presence of the loxP sites, while PCR-3 confirmed the presence of the *Stra8-Cre* transgene. *represents internal control PCR bands.

**(C)** Western blotting shows analysis the abundance of SMARCA5 protein in the testes from *Cre/+* and *Smarca5 fl/fl* mice at 14 and 30 days postpartum (dpp). ACTIN served as a loading control. Each lane contained 75 μg total testis protein extract.

**(D)** Immunofluorescence staining for SMARCA5 in the Bouin-fixed testis sections from *Cre/+* and *Smarca5-CKO* mice at 30 dpp. S, Sertoli cells; Z, zygotene spermatocytes; P, pachytene spermatocytes. Bars represent the quantification of SMARCA5-positive cells per seminiferous tubule from three independent biological replicates (*n* = 3). Whiskers show standard deviation, and the center of the whiskers represents the mean. Significance is measured by two-sided unpaired Student *t* test. *p* = 4.4 × 10^–13^. Scale bars: 50 μm.

**(E)** Immunofluorescence staining for SOX9 in the Bouin-fixed testis sections from *Cre/+* and *Smarca5-CKO* mice at 14 and 30 dpp. Insets show enlarged views of the boxed regions. Bars represent the quantification of SOX9-positive Sertoli cells per seminiferous tubule from three independent biological replicates (*n* = 3). Whiskers show standard deviation, and the center of the whiskers represents the mean. Significance is measured by two-sided unpaired Student *t* test. N.s. not significant. Scale bars: 50 μm.

Figure S4. ****Germ cell-specific deletion of Smarca5 causes reduced testis size and the absence of epididymal spermatozoa. Related to Figure 2.****

**(A)** Comparison of body size and testes from 8-week-old *Cre/+* and *Smarca5-CKO* mice. Representative images of right (R) and left (L) testes are shown.

**(B)** Relative testes weights from *Cre/+* and *Smarca5-CKO* mice at 14 days postpartum (dpp) and 30 dpp (*n* = 3). Significance is measured by two-sided unpaired Student *t* test.

**(C)** Hematoxylin and eosin staining of caudal epididymis sections from 8-week-old *Cre/+* and *Smarca5-CKO* mice. Scale bars: 50 μm.

Figure S5. ****SMARCA5-deficient germ cells fail to progress through prophase I of meiosis. Related to Figure 2.****

**(A–D)** Immunofluorescence staining of the testis sections from *Cre/+* and *Smarca5-CKO* mice at 14 and 30 days postpartum (dpp) for **(A)** DDX4 (14 dpp, *n* = 3, *p* = 1.6 × 10^–5^; 30 dpp, *n* = 3, *p* = 1.1 × 10^–12^), **(B)** TUNEL (14 dpp, *n* = 3, *p* = 9.1 × 10^–6^; 30 dpp, *n* = 3, *p* = 7.2 × 10^–7^), **(C)** HSPA2 (30 dpp, *n* = 3, *p* = 8,5 × 10^–15^), and **(D)** PNA (30 dpp, *n* = 3, *p* = 5.4 × 10^–17^). Insets show higher magnification views of the boxed regions. Bars represent the quantification of DDX4-, TUNEL-, HSPA2-, and PNA-positive germ cells per seminiferous tubule from three independent biological replicates. Whiskers show standard deviation, and the center of the whiskers represents the mean. Significance is measured by two-sided unpaired Student *t* test. Scale bars: 50 μm.

**Figure S6. The genome-wide SMARCA5 and BPTF occupancies are strongly correlated across biological replicates. Related to Figure 4.**

**(A)** Heatmap shows the Pearson correlation coefficient (*r*) between the biological replicates of SMARCA5 CUT&Tag biological replicates from spermatogonia (SpG).

**(B)** Heatmap shows the Pearson correlation coefficient (*r*) between the biological replicates of BPTF CUT&Tag biological replicates from spermatogonia (SpG).

**Figure S7. The Table for the known HOMER motif analysis at the promoters of upregulated genes *in Smarca5-CKO* mice. Related to Figure 4.**

**Figure S8. Promoter occupancy patterns of NFYA, SMARCA5, and BPTF at non-poised meiosis I and spermiogenesis genes in spermatogonia. Related to Figure 5.**

Heatmaps show the reads per million (RPM)-normalized ATAC-seq signal (replicates 2 and 3), together with NFYA, SMARCA5, and BPTF occupancy (replicates 2 and 3), in the −5 kb to +5 kb regions flanking the transcription start sites (TSSs) and transcription end sites (TESs) of poised and non-poised genes in spermatogonia (SpG). Genes are ordered according to ATAC-seq signal intensity.
