## Supplementary figures and images for "Active maintenance of accessible chromatin at poised meiotic promoters by the NFYA-NURF axis is required for meiotic entry in mice"

### Figure S1

Eghbali et al., Figure S1.

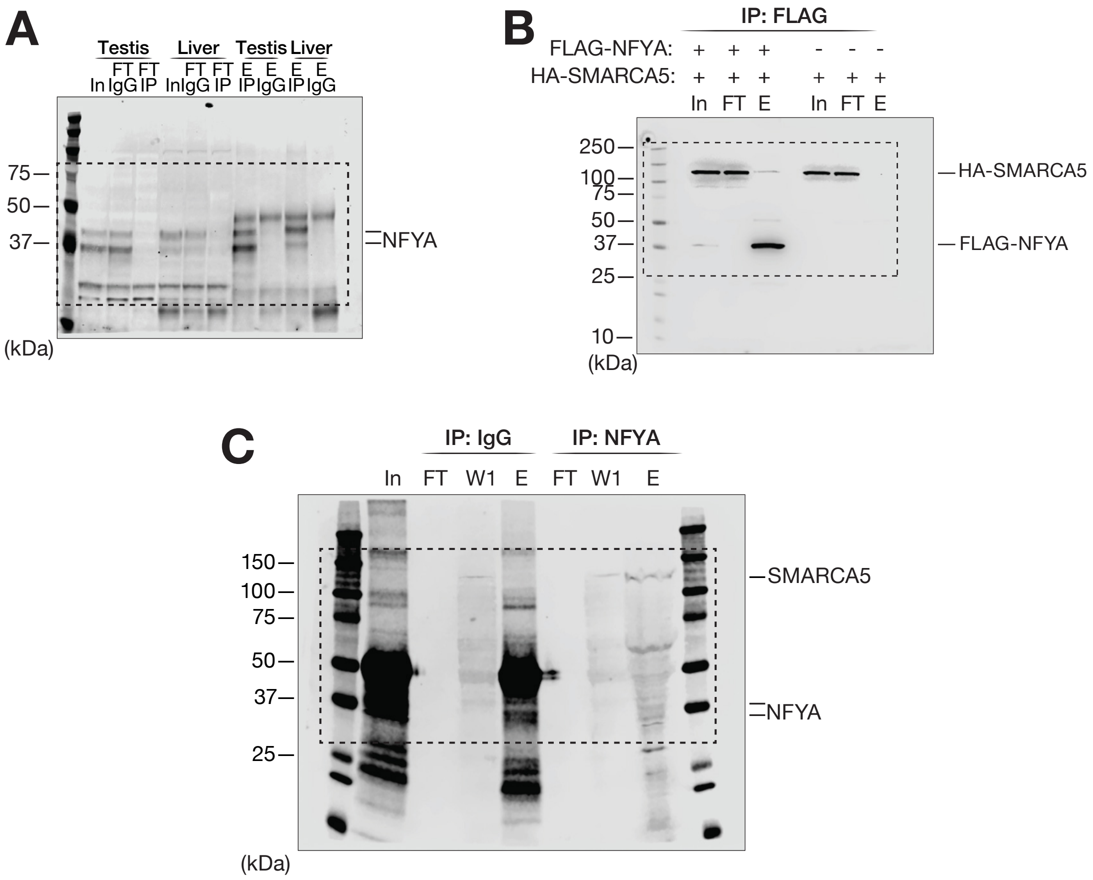

### Figure S2

Eghbali et al., Figure S2.

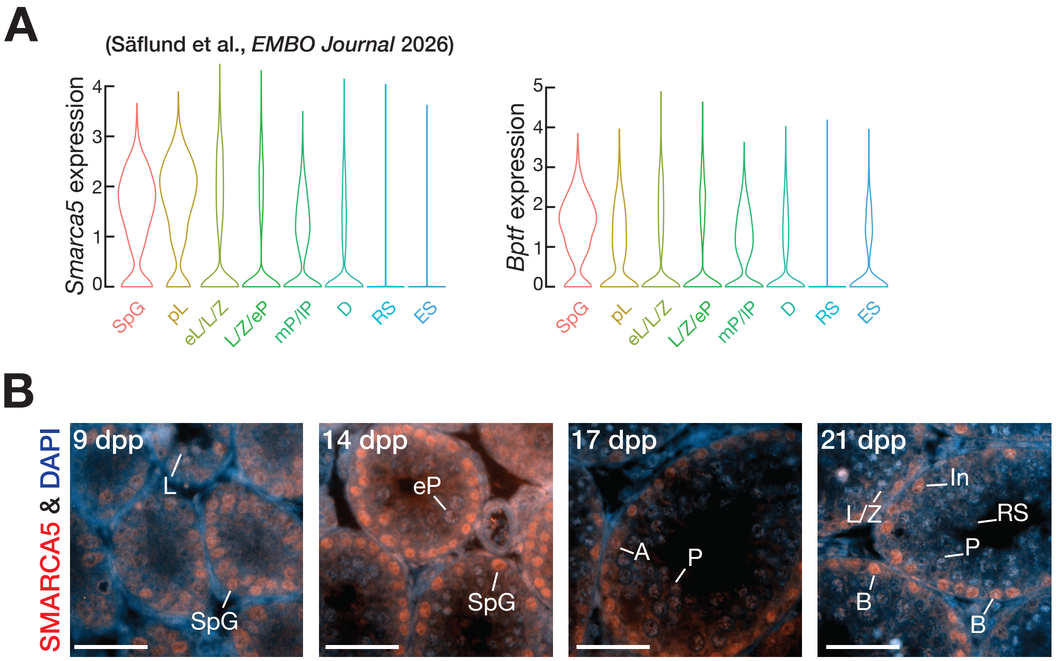

### Figure S3

Eghbali et al., Figure S3.

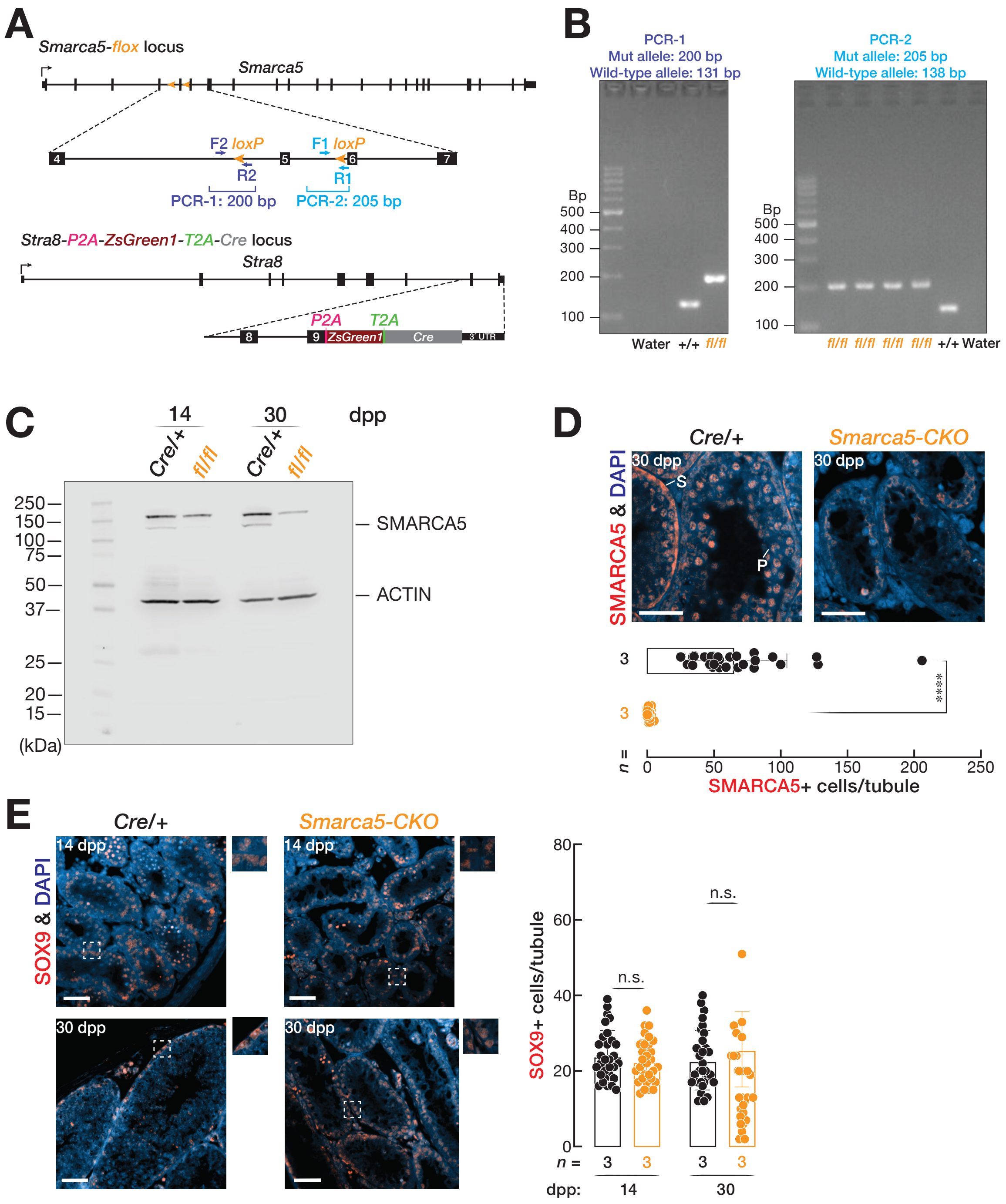

### Figure S4

Eghbali et al., Figure S4.

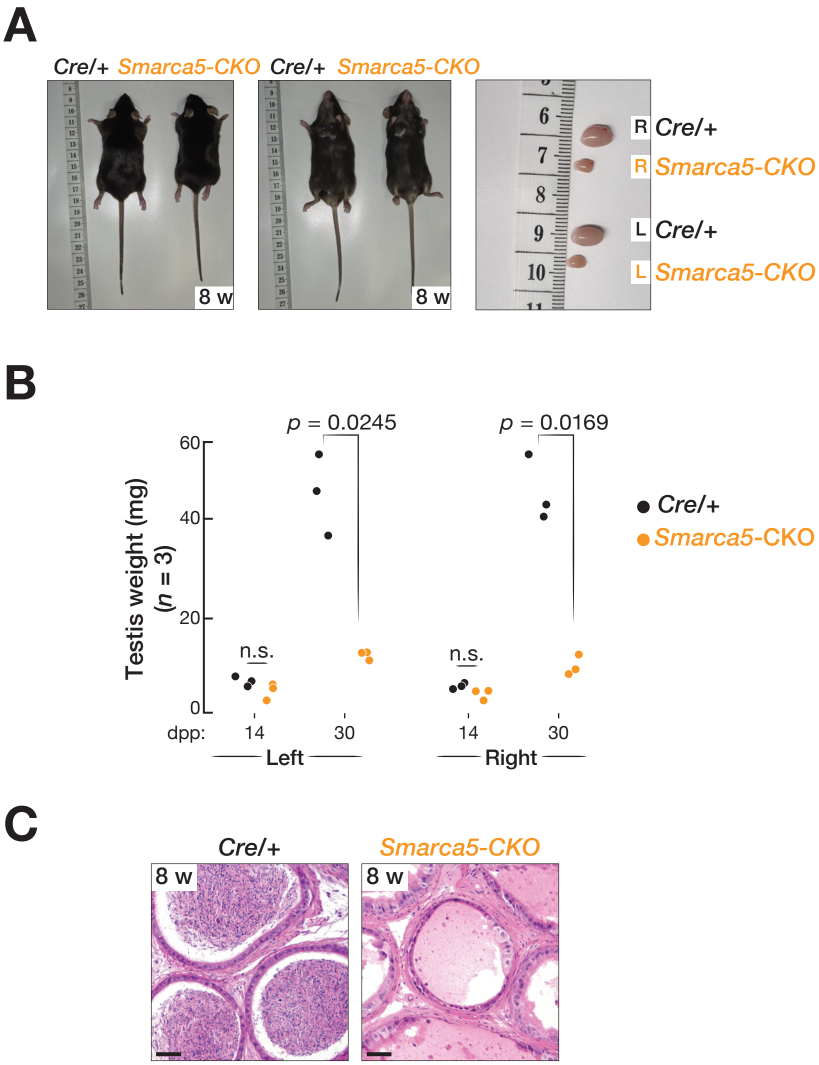

### Figure S5

Eghbali et al., Figure S5.

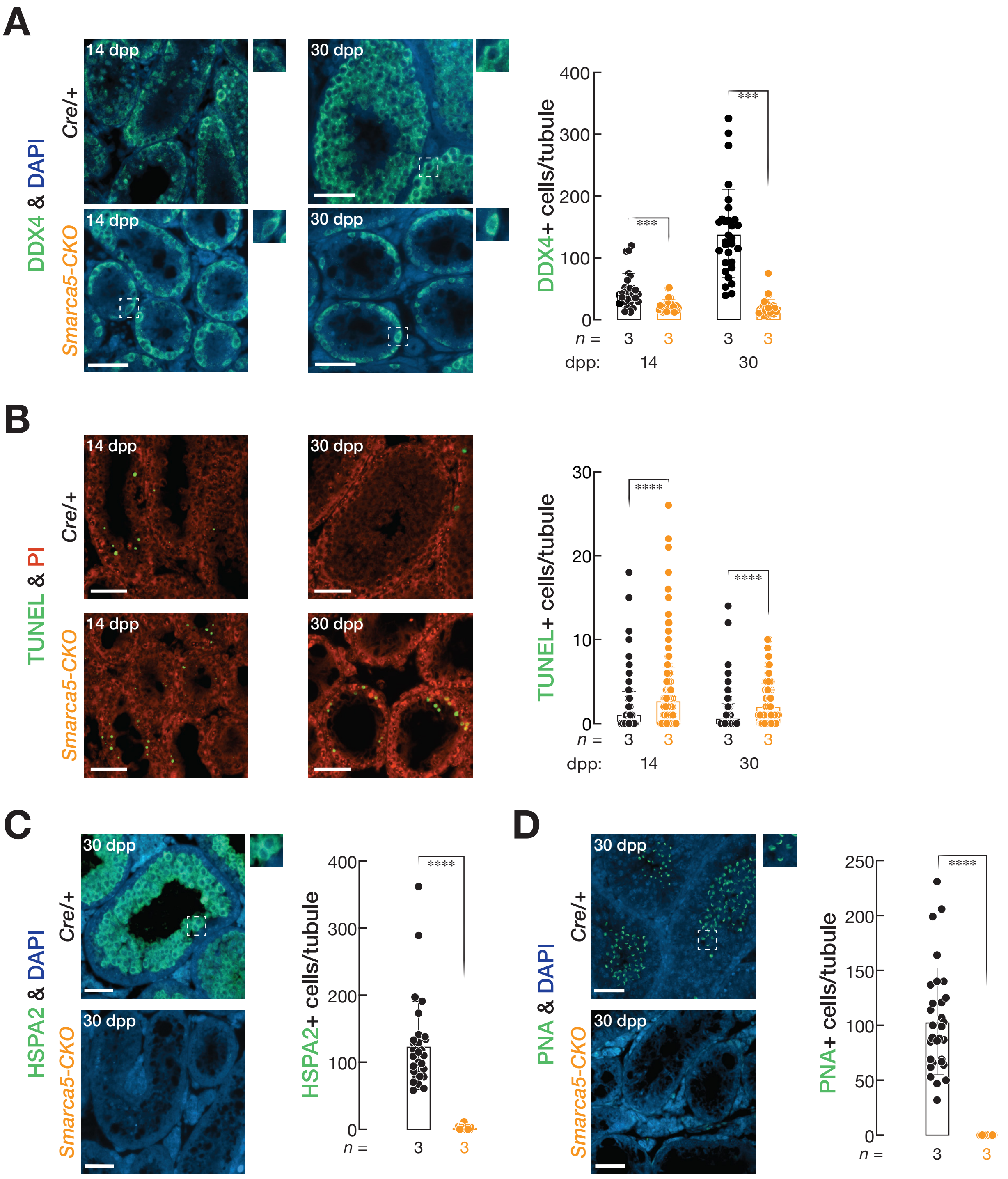

### Figure S6

Eghbali et al., Figure S6.

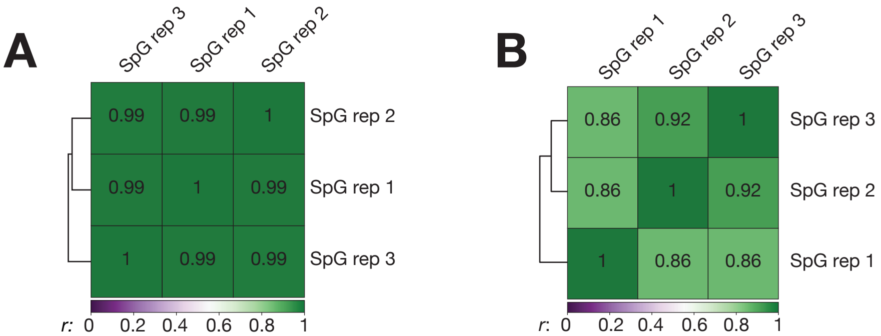

### Figure S8

Eghbali et al., Figure S8.

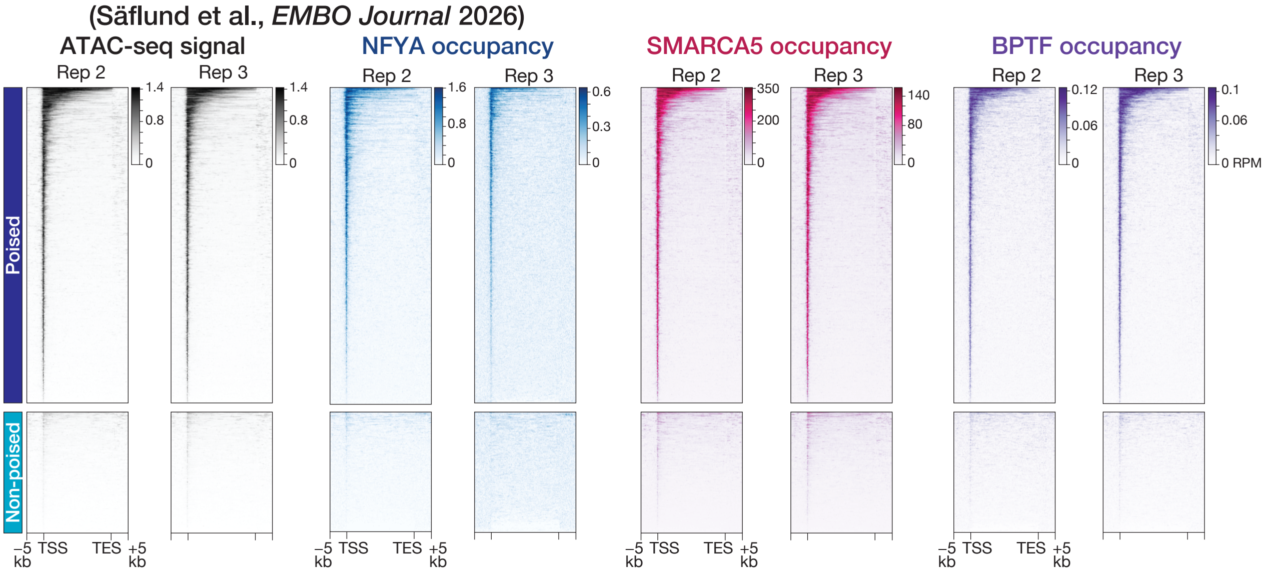
