## Supplementary material for "Active maintenance of accessible chromatin at poised meiotic promoters by the NFYA-NURF axis is required for meiotic entry in mice": Figure S7

Eghbali et al., Figure S7.

### Known motifs at increased gene promoters from top 25 GO terms

| Rank | Motif | Factor | *p* value | Change in RNA expression of the factor SpG_*Smarca5-CKO* ÷ SpG *Cre*/+ |
|---|---|---|---|---|
| 1 | 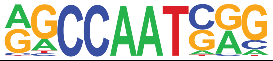 | NFY | $10^{-5}$ | — |
| 2 | 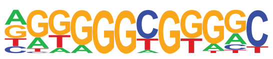 | KLF15 | $10^{-5}$ | ↑ |
| 3 | 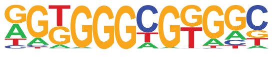 | KLF4 | $10^{-5}$ | — |
| 4 | 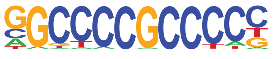 | SP1 | $10^{-4}$ | — |
| 5 | 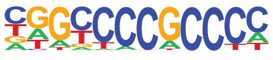 | SP2 | $10^{-4}$ | — |
| 6 | 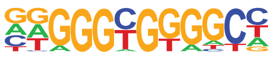 | KLF1 | $10^{-4}$ | — |
| 7 | 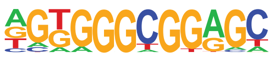 | SP5 | $10^{-3}$ | ↑ |
| 8 | 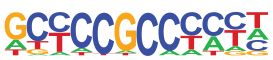 | KLF17 | $10^{-3}$ | ↓ |
| 9 | 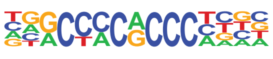 | KLF3 | $10^{-3}$ | ↑ |
| 10 | 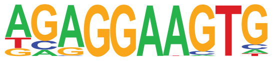 | PU.1 | $10^{-3}$ | — |
| 11 | 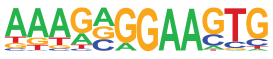 | SPIB | $10^{-3}$ | — |
| 12 | 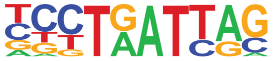 | LHX6 | $10^{-3}$ | — |
| 13 | 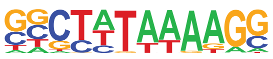 | TFIID | $10^{-3}$ | — |
